# Parvalbumin-expressing basal forebrain neurons mediate learning from negative experience

**DOI:** 10.1101/2023.03.31.535018

**Authors:** Panna Hegedüs, Victoria Lyakhova, Anna Velencei, Márton I. Mayer, Zsofia Zelenak, Gábor Nyiri, Balázs Hangya

## Abstract

Parvalbumin (PV)-expressing GABAergic neurons of the basal forebrain (BFPVNs) were proposed to serve as a rapid and transient arousal system. While they have a well-documented role in the regulation of sleep-wake states, whether and how BFPVNs participate in mediating awake behaviors is not known. To address this, we performed bulk calcium imaging and recorded single neuronal activity from the horizontal band of the diagonal band of Broca (HDB) while mice were performing an associative learning task. Genetically identified BFPVNs of the HDB responded with a distinctive, phasic activation to punishment. In contrast, reward only elicited slow and delayed responses, while stimuli predicting behavioral reinforcement (reward or punishment) were followed by a gradual increase of HDB BFPVN firing rates. Optogenetic inhibition of HDB BFPVNs during punishment impaired the formation of cue-outcome associations, suggesting a causal role of these neurons in associative learning. Mapping the input-output connectivity of HDB BFPVNs by anterograde and mono-transsynaptic retrograde tracing experiments showed that these neurons received strong inputs from the hypothalamus, the septal complex and the median raphe region, while they synapsed on diverse cell types in key structures of the limbic system including the medial septum, the retrosplenial cortex and the hippocampus. Bulk calcium imaging performed in these termination regions indicated that HDB BFPVNs broadcast information about aversive stimuli to multiple downstream targets. We propose that the arousing effect of BFPVNs is recruited by aversive stimuli to serve crucial associative learning functions during awake behaviors.

## Introduction

Basal forebrain nuclei in the ventral part of the forebrain are defined by the presence of cholinergic projection neurons (BFCNs)^1–3^. However, there are approximately five times more GABAergic than cholinergic neurons in the BF^4–6^, of which the parvalbumin-expressing population was shown to project to several cortical and subcortical target areas^7–9^.

These BFPVNs were shown to be important regulators of cortical gamma oscillations^10^ and to promote wakefulness and arousal through their local^11^ and distant projections^12^. However, a number of observations point to the direction that BFPVNs might also play a role in awake behaviors, and associative learning in particular. First, similarly to BFCNs, degeneration of BFPVNs was also linked to cognitive decline in Alzheimer’s disease^13^. Second, non-selective lesions of the BF revealed a more substantial attentional and learning deficit in rodents compared to selective cholinergic lesions^14–17^ and a study of selective BF GABAergic ablation revealed a deficit in eyeblink conditioning^18^. Third, a line of studies demonstrated that non-cholinergic BF neurons respond phasically to behaviorally salient stimuli irrespective of their valence^19^, where stronger responses^20^ or higher pre-stimulus activity^21^ predicted faster and more precise decision speed during operant conditioning. These studies raise the possibility, that BFPVNs might also play a role in associative learning.

To address this question, we used in vivo electrophysiological recording with optogenetic tagging and bulk calcium imaging by fiber photometry of PV-expressing neurons of the rostral HDB nucleus of the BF (referred to as BFPVNs hereinafter), while mice were performing a probabilistic Pavlovian conditioning task^22^. We found that BFPVNs showed rapidly rising, brief (i.e., ‘phasic’) responses to punishment, whereas rewards, as well as sensory stimuli that predicted reward or punishment, elicited comparably smaller and delayed responses. These observations revealed that BFPVN activity correlated with behavioral reinforcement, and punishment in particular, suggesting that these neurons signaled negative behavioral outcome to other parts of the brain. In accordance, optogenetic inhibition of BFPVNs during punishments in a Pavlovian conditioning task impaired the ability of mice to form stimulus-outcome associations, demonstrating that HDB BFPVNs encode outcome-related information crucial for associative learning.

We performed cell type-specific anterograde and mono-transsynaptic retrograde tracing experiments to reveal long range afferent and efferent connections of HDB BFPVNs. We demonstrated that they received dense inputs from the lateral hypothalamus, the septal complex and the median raphe nucleus, and innervated important nodes of the limbic system crucial for contextual learning, including the medial septum, the hippocampal formation and the retrosplenial cortex^23–28^. They synapsed on a broad range of cell types including cholinergic and PV-expressing neurons in the BF, as well as interneuron types of the neocortex and hippocampus, as revealed by light- and electron microscopy. Finally, by performing bulk calcium imaging of HDB BFPVN fibers in target areas, we demonstrated that these neurons broadcast outcome-related information to multiple downstream areas. These data suggest that BFPVNs of the HDB mediate learning from negative reinforcement.

## Results

### Identified BFPVNs of the HDB show phasic responses to punishment

We trained PV-Cre mice (n = 4) on an auditory probabilistic Pavlovian conditioning task (Hegedüs et al., 2021) to examine how HDB BFPVNs respond to cues predicting reward or punishment with different probabilities and to the reinforcement itself. Mice were head restrained and two tones of different pitch were presented in a randomized order (Fig.1a); Cue 1 predicted reward with higher probability (water drop; 80% reward, 10% punishment, 10% omission), whereas Cue 2 predicted punishment with higher probability (air puff; 65% punishment, 25% reward, 10% omission). This experimental design enabled us to examine how BFPVNs respond to surprising and expected reward or punishment (unconditioned stimuli, US) as well as their predicting cues (conditioned stimuli, CS). Mice learned these task contingencies, demonstrated by significantly higher anticipatory lick rate in response to Cue 1 that predicted likely reward (Fig.1b-e).

To examine how individual BFPVNs responded to conditioned and unconditioned stimuli, we injected AAV2.5-EF1a-DiO-hChR2(H134R)-eYFP.WPRE.hGh into the HDB region of PV- Cre mice and performed extracellular multiple single-cell recordings combined with optogenetic tagging of HDB BFPVNs (n = 36) by implanting a head-mounted micro-drive housing moveable tetrode electrodes and an optical fiber (Fig.1f-j, Fig.S1; specific expression verified in ref. ^29^, Figure S15). Electrode placement in the HDB was verified by histological reconstruction at the end of the experiment (Fig.1f-g). We found that BFPVNs responded with a gradual, ramp-like activation to cue onset (n = 12/36, 33%; Fig.2a-b) irrespective of the predictive value of the cues (Fig.S2) and with a small but significant activation to the delivery of water reward (n = 14/36, 39%; Fig.2c-d). In contrast, they responded to punishment with fast, phasic activation, homogeneously detectable in most (n = 26/36, 72%) identified BFPVNs (Fig.2e-f). BFPVN punishment responses showed adaptation, as they decreased in magnitude as a function of repeated air-puff presentations; however, they remained present in late trials of the training sessions (Fig.2g-i). BFPVN responses to reward and punishment were not modulated by surprise (Fig.S2), unlike what we found in case of reward-elicited responses of BFCNs in the same task^30, 31^.

We examined the electrophysiological properties of punishment responsive BFPVNs and found that half of these neurons were burst firing. Both burst spikes and single spikes were time-locked to punishment presentation, but bursting BFPVNs were more punishment-activated (n = 17/18, 94%) than non-bursters (n = 10/18, 56%; Fig.S3).

### Optogenetic inhibition of HDB BFPVNs during punishment impairs association learning

To assess whether the robust responses of BFPVNs to punishment represent an aversive signal inducing avoidance behavior, we tested whether optogenetic activation of BFPVNs elicited conditioned place aversion. We injected either AAV2.5-EF1a-DiO-hCHR(H134R)- eYFP.WPRE.hGh (n = 8) or AAV2.5-EF1a-DiO-eYFP.WPRE.hGh (n = 10 used as controls) viral vectors into the HDB of PV-Cre mice and performed a conditioned place aversion test. Mice were placed in an arena, the two sides of which were separated by a small open gate and the sides were differentiated by dotty and striped sidewall patterns. After habituation, BFPVNs were optogenetically stimulated in one side of the arena and different behavioral parameters such as total distance traveled, speed, time spent in either side of the arena, grooming, digging, freezing, and rearing were recorded and analyzed. We found that stimulation of HDB BFPVNs did not elicit aversive (or appetitive) behavior, as channelrhodopsin-expressing mice did not avoid the stimulated side, nor did they express any other behavioral differences compared to control animals, and they did not show behavioral signs of stress (Fig.S4).

Next, to causally test the behavioral function of the punishment-elicited HDB BFPVN response in learning, we tested whether the optogenetic suppression of BFPVNs during the time of air-puff delivery affected the learning ability of mice. We injected the HDB of PV-Cre mice bilaterally with AAV2.9-CAG-Flex-ArchT-GFP (n = 6, ArchT group) or AAV2.9-CAG-Flex - GFP (n = 8, control group; Fig.3a). A continuous laser pulse was applied to HDB BFPVNs through bilaterally implanted optic fibers upon punishment presentation while mice were trained on the auditory Pavlovian conditioning task. Both ArchT-expressing and control mice developed anticipatory licking in response to the conditioned stimuli. However, while control mice learned to suppress licking after Cue 2 that signaled likely punishment (similar to our initial cohort, see Fig. 1b-e), lick rates of ArchT-expressing mice remained high after both cues and thus they did not show differential anticipatory licking after the CS (Fig.3b-c). Receiver operating characteristic analysis (ROC; see Methods) revealed a significant difference between ArchT-expressing and control mice in a half-second window before reinforcement delivery (from about 0.6 to 1.1 s after cue onset, Fig.3d, Fig.S5). Control mice not only responded with a higher anticipatory lick rate to Cue 1 predicting likely reward, but their responses were significantly faster compared to those after Cue 2 predicting likely punishment. This reaction time difference was also abolished by suppressing HDB BFPVNs in the ArchT group of mice (Fig.3e-f). Thus, optogenetic suppression of HDB BFPVNs prevented mice from associating higher value with the cue that predicted likely reward.

**Figure 1.**
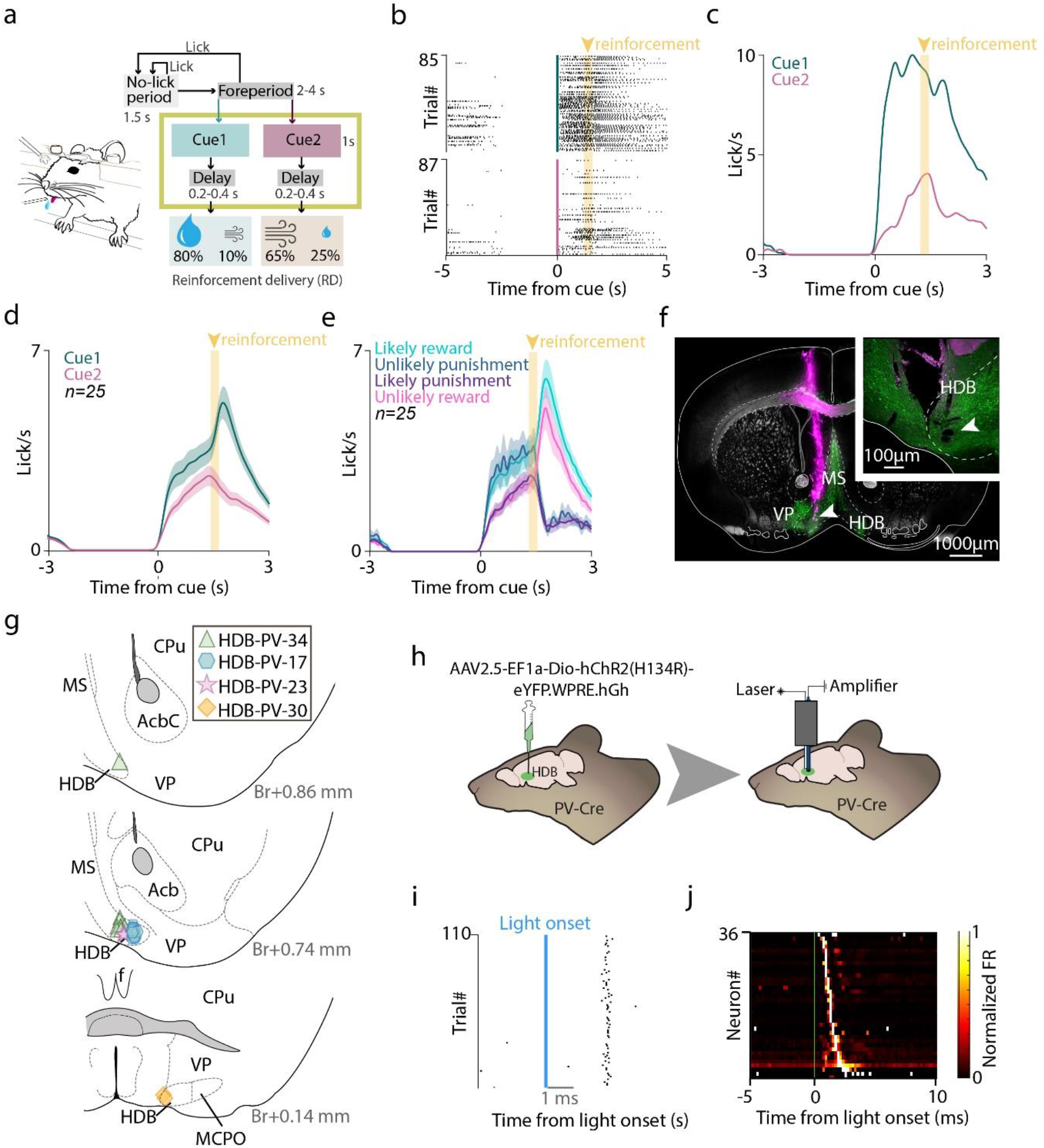
Head-fixed probabilistic Pavlovian conditioning combined with tetrode recording and optogenetic tagging. **a,** Schematic diagram of the task protocol. After a variable foreperiod, in which the animals were not allowed to lick, they were presented with two cues, either predicting likely reward or likely punishment; reinforcement was preceded by a short, variable delay period. **b,** Raster plot of mouse licking times aligned to stimulus (cue) onset from an example session, showing higher anticipatory lick rate in response to the reward-predicting cue (Cue 1). **c,** Lick rates (peri-event time histograms, PETH) aligned to stimulus onset from the same example session. **d,** Average lick rates of all recording sessions in which BFPVNs were identified (n = 25 sessions). **e,** Average lick rates of the same sessions as in D, partitioned according to the four possible outcomes. **f,** Histological reconstruction of the tetrode track in the HDB. Green, ChR2-eYFP; magenta, DiI. Top right, magnified image of the electrolytic lesion site (white arrowhead), indicating the tip of the electrodes. MS, medial septum; VP, ventral pallidum. **g,** Locations of the recorded BFPVNs based on histological reconstruction. Different markers correspond to different mice. Numbers show antero-posterior positions relative to Bregma. **h,** Schematic illustration of viral injections, extracellular tetrode recordings and optogenetic tagging in mice. **i,** Example raster plot of spike times aligned to photostimulation pulses of an optogenetically identified BFPVN. **j,** Firing rate (color-coded PETH) of all optogenetically identified BFPVNs aligned to photostimulation onset.

**Figure 2.**
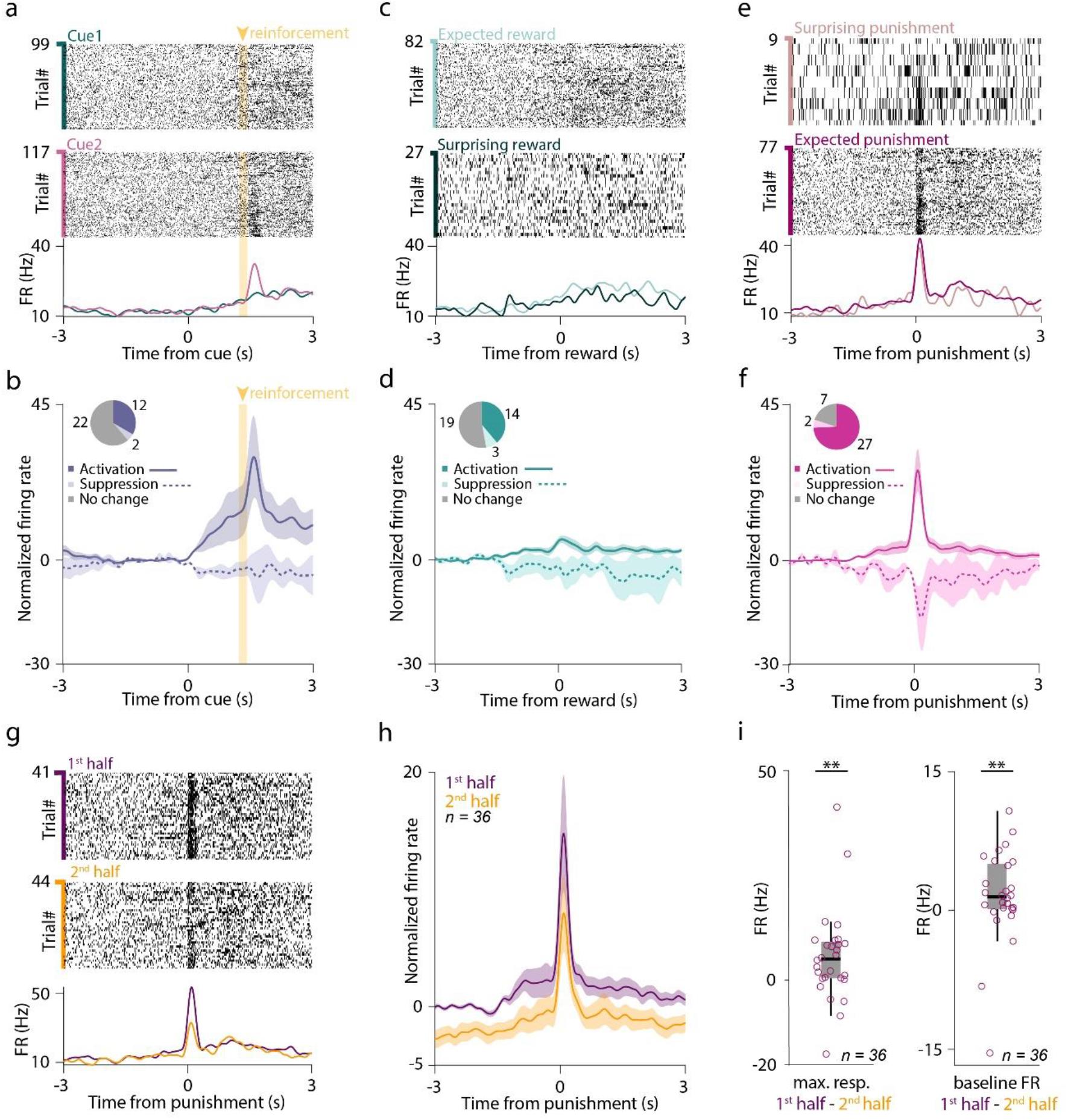
BFPVNs of the HDB show strong responses to punishment. **a,** Raster plots and PETHs of the spike times aligned to cue onset of an example BFPVN. Trials are partitioned by the cue types (Cue 1 predicting likely reward / unlikely punishment, top, green; Cue 2 predicting likely punishment / unlikely reward, middle, purple). **b,** Average PETH of BFPVN activity aligned to cue onset (n = 36). **c,** Example PETH of the same BFPVN as in panel A aligned to reward delivery. Trials are partitioned to expected (following Cue 1) and surprising (following Cue 2) reward trials. **d,** Average PETH of BFPVN activity aligned to reward delivery (n = 36). **e,** Example PETH of the same BFPVN as in panel A aligned to punishment delivery. Trials are partitioned to surprising (following Cue 1) and expected (following Cue 2) punishment trials. **f,** Average PETH of BFPVN activity aligned to punishment delivery (n = 36). **g,** Spike raster plots and PETH aligned to punishment delivery for an example BFPVN. Top, trials in the first half of the sessions; middle, trials in the second half of the session; bottom, PETH. **h,** Average PETH of BFPVN activity aligned to punishment delivery (n = 36). Purple, first half of the session; mustard, second half of the session. **i,** Left, difference in peak punishment response between the first and second half of the session (**, p < 0.01, p = 0.009108, Wilcoxon signed- rank rest). Right, difference in baseline firing rate between the first and second half of the session (**, p < 0.01, p = 0.00135, Wilcoxon signed-rank test).

**Figure 3.**
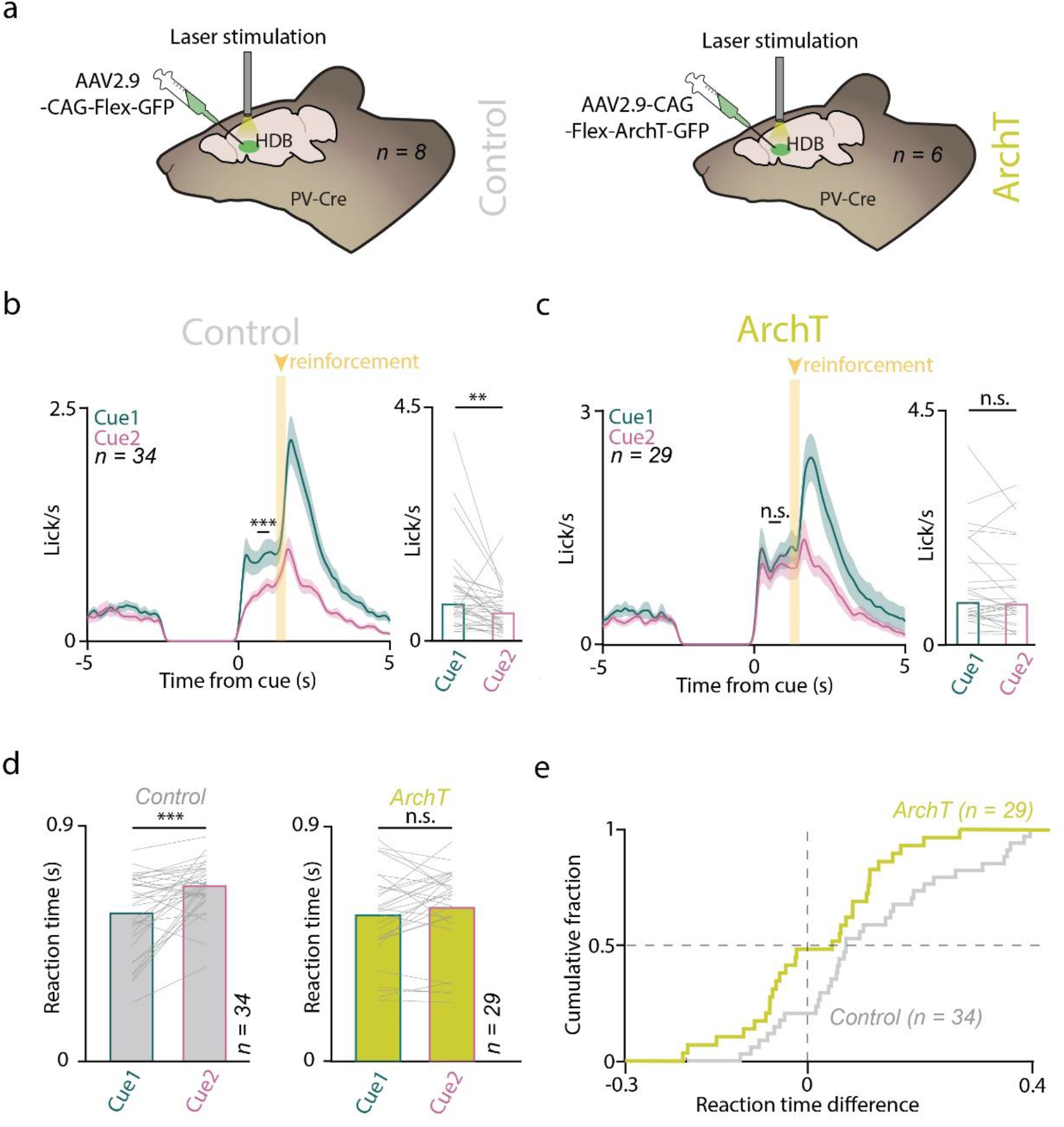
Optogenetic suppression of HDB BFPVN punishment responses disrupts learning cue contingencies during Pavlovian conditioning. **a,** Schematic illustration of optogenetic suppression of BFPVNs in PV-Cre mice. **b,** Left, average lick rate of control mice in the final training stage of the task (n = 34 sessions). ***, p < 0.001, ROC analysis with permutation test (see Methods). Right, bar graph showing mean lick rate before reinforcement (0.6-1.1 s window from cue onset, see ROC analysis in Fig.S4). Lines correspond to individual sessions. **, p < 0.01, p = 0.001517, Wilcoxon signed-rank test. **c,** Left, average lick rate of ArchT-expressing mice in the final training stage of the task (n = 29 sessions). n.s., p > 0.05, ROC analysis. Right, bar graph showing mean lick rate in a 0.6-1.1 s window from cue onset. n.s., p > 0.05, p = 0.40513, Wilcoxon signed-rank test. **d,** Left, reaction time difference between lick responses to Cue 1 and Cue 2 in control animals. ***, p < 0.001, p = 0.000472, Wilcoxon signed-rank test. Right, same for the ArchT group. n.s., p > 0.05, p = 0.15668, Wilcoxon signed- rank test. **e,** Cumulative histogram of reaction time differences (Cue 2 – Cue 1) of ArchT (yellow, n = 29 sessions) and control (grey, n = 34 sessions) animals.

### BFPVNs of the HDB receive inputs from aversion-coding nuclei and send projections to the limbic system

We used mono-transsynaptic rabies tracing to identify brain areas and cell types that provide synaptic input to HDB BFPVNs. We injected Cre-dependent helper virus and G protein-deleted EnvA-pseudotyped rabies virus into the HDB of PV-Cre mice (n = 3; Fig.4). Long-range input cells were distributed throughout the whole brain, suggesting a high level of input convergence onto HDB BFPVNs (Table S1.). The hypothalamus provided the most inputs (lateral hypothalamus, 33.1%, fraction of total input neurons; preoptic area, 7.6%), followed by the lateral septum (13.8%). Neighboring areas of the basal forebrain, such as the medial septum (MS, 10.8%) and the vertical limb of the diagonal band of Broca (VDB, 6.4%) also contained substantial number of input cells, together with the nucleus accumbens (3.5%), while the median raphe region (MRR) provided the highest number of synaptic inputs from the brainstem (3.9%, Fig.4e). In case of one mouse, we performed unilateral helper and rabies virus injection to the HDB. In this animal, the presynaptic neurons were predominantly ipsilateral (93.7%) to the starter population, and the pattern of anatomical connectivity was similar for ipsilateral and contralateral projections.

**Figure 4.**
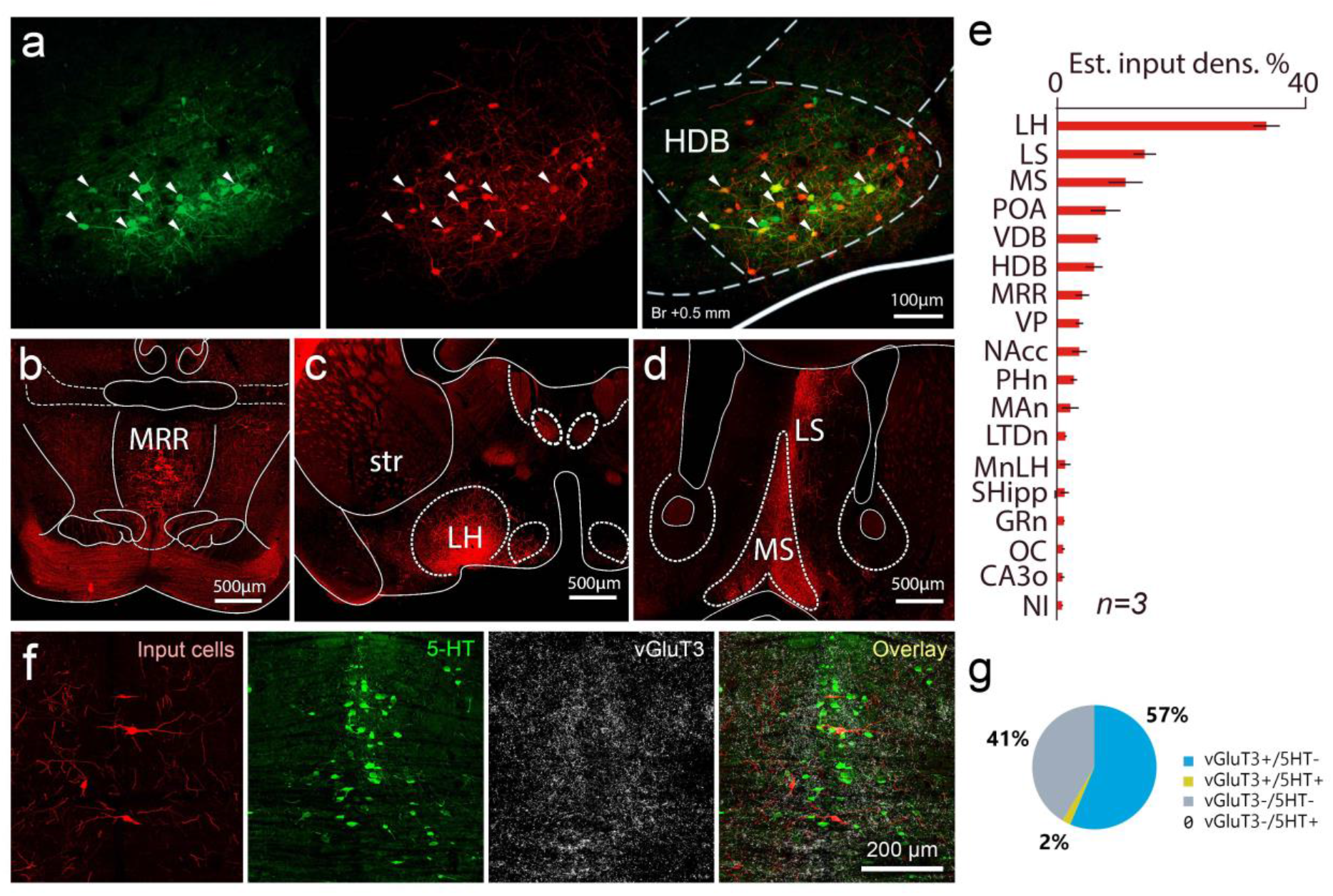
BFPVNs of the HDB receive inputs from aversion-coding nuclei. **a,** Retrograde tracing, using pseudotyped rabies virus. Green, helper virus (left); red, pseudotyped rabies virus (middle); yellow, colocalizaton (right). Starter cells in the HDB are indicated by white arrowheads. **b,** Coronal fluoromicrograph showing input cells in the MRR (red fluorescence). **c,** Coronal fluoromicrograph showing input cells in the lateral hypothalamus (red fluorescence). **d,** Coronal fluoromicrograph showing input cells in the medial septum (red fluorescence). **e,** Estimated input density (% of all input cells) in different input brain regions. **f,** Immunohistochemical staining of input neurons (red), serotonergic neurons (5-HT, green) and vGluT3+ glutamatergic neurons (vGluT3, far red visualized in white) in the MRR. **g,** Proportion of neurons in the MRR that target BFPVNs. Abbreviations: LH, lateral hypothalamus; LS, lateral septum; MS, medial septum; POA, preoptic area; VDB; vertical limb of the diagonal band of Broca; HDB, horizontal limb of the diagonal band of Broca; MRR, median raphe region; VP, ventral pallidum; NAcc, nucleus accumbens; PHn, posterior hypothalamic nucleus; MAn, medial amygdaloid nucleus; LTDn, laterodorsal tegmental nucleus; MnLH, magnocellular nucleus of the lateral hypothalamus; SHipp, septohippocampal nucleus; GRn, gigantocellular reticular nucleus; OC, orbital cortex; CA3o, CA3 stratum oriens; NI, nucleus incertus.

Because MRR has an important role in mediating negative experience, we further investigated its cell types that provided inputs to HDB BFPVNs. MRR vesicular glutamate transporter 2 (vGluT2) expressing cells that are key regulators of negative experience are known to target BFPVNs^32^. However, MRR has several other types of cells including serotonergic (5HT) neurons, vGluT3-expressing glutamatergic neurons, those that express both serotonin and vGluT3, and GABAergic neurons as well^33^. We found that most of the input cells of BFPVNs were vGluT3-expressing glutamatergic neurons (59%, n = 2 mice) and out of these cells only a small fraction were serotonergic as well (5-HT-immunopositive, 2% of all inputs from MRR, Fig.4f-g). We found no input cells that were only 5HT-immunopositive. The rest (41%) of the input cells (that were immunonegative for both vGluT3 and 5-HT) included vGluT2 cells that are known to target BFPVNs and may also include some MRR GABAergic neurons. These suggest that MRR targets BFPVNs with both vGluT2- and vGluT3-expressing glutamatergic neurons, both of which play a key role in the brainstem control of negative experience^32, 34^.

Next, we examined the downstream targets of HDB BFPVNs by performing a series of anterograde tracing experiments. We injected AAV2.9-CAG-Flex-eGFP viral vectors bilaterally in the HDB of PV-Cre mice to visualize HDB BFPVN axons using fluorescent microscopy (n = 3; Fig.5a). We found that the majority of the projecting axons targeted the medial septum – diagonal band of Broca area, the hippocampus and the retrosplenial cortex (Fig.5a-b), which are important regions of the limbic system. Additionally, other limbic structures such as the medial mammillary nucleus, the supramammillary nucleus and the paratenial thalamus were also targeted by HDB BFPVNs. A smaller extent of projections was received by the prefrontal cortex, the lateral septum and the MRR. By using correlated electron and light microscopy, we demonstrated that HDB BFPVNs established symmetrical synaptic contacts with PV-expressing and cholinergic neurons in the medial septum, with calretinin-expressing (CR+) and PV-expressing neurons in the hippocampal CA1 region and with PV- expressing neurons in the retrosplenial cortex (Fig.5b-d).

**Figure 5.**
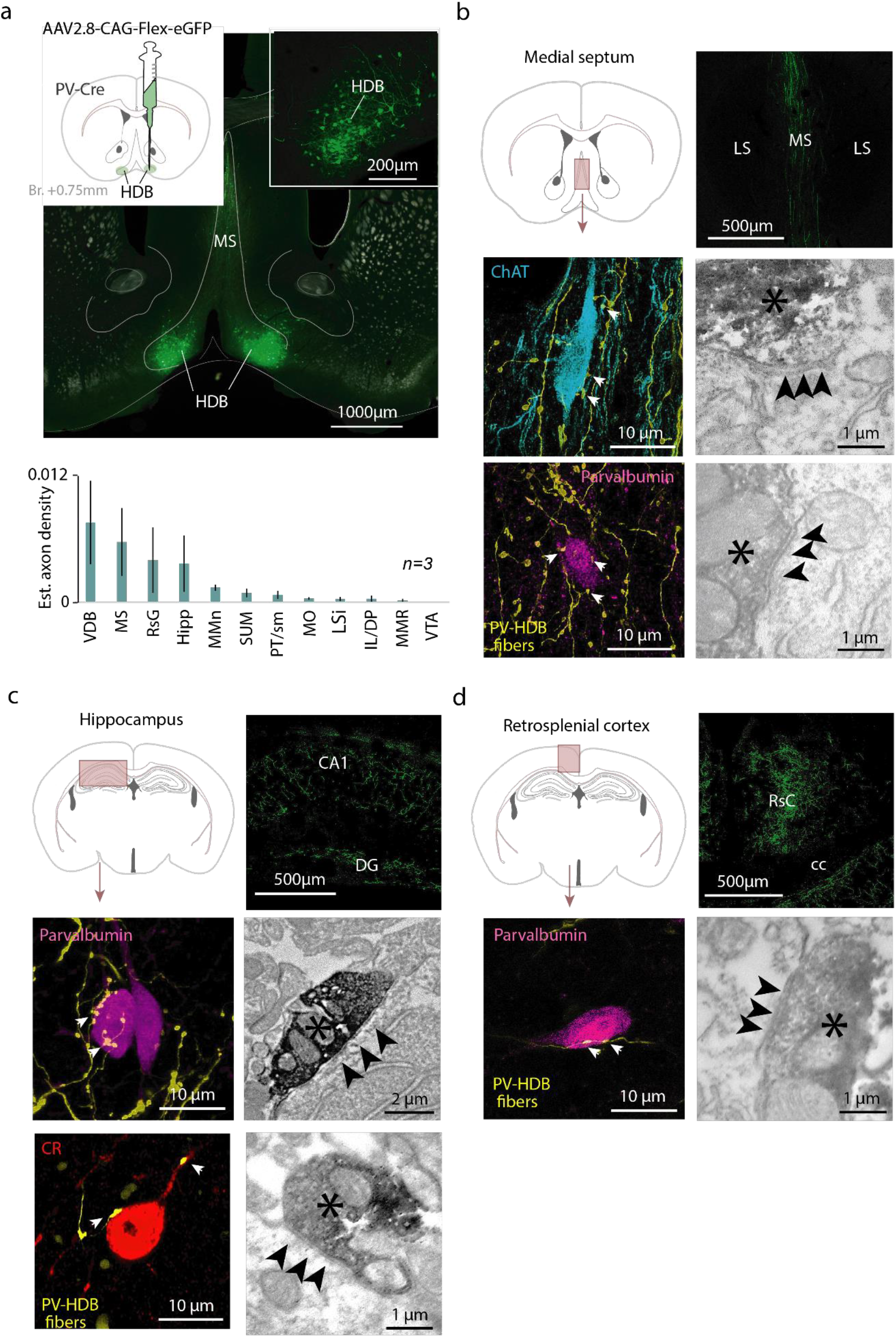
BFPVNs of the HDB innervate the limbic system. **a,** Top left, schematic illustration of cell-type specific anterograde tracer virus injection into the HDB. Bottom left, fluorescent image of the injection site (green, eGFP). Right, normalized BFPVN axon density in targes brain areas (median ± SE of median). **b,** Top left, schematic illustration of the medial septum on a coronal atlas image. Top right, fluoromicrograph of labelled axons in the MS. Middle and bottom left, fluorescent images of ChAT+ and PV+ target neurons in the medial septum (white arrowheads indicate plausible contact sites). Right, electronmicrographs of synaptic contacts established by BFPVNs with the target cell types shown in the middle panels (different examples). HDB BFPVN axons are labelled by DAB precipitate (black asterisk); black arrows indicate the synaptic contacts. **c,** Top left, schematic illustration of the hippocampus on a coronal atlas image. Top right, fluoromicrograph of labelled axons in the CA1. Middle and bottom left, fluorescent images of CR+ and PV+ target neurons in the hippocampus. Right, electronmicrographs of synaptic contacts established by BFPVNs with the targes cell types shown in the middle panels. **d,** Top left, schematic illustration of the retrosplenial cortex (RsC) on a coronal atlas image. Top right, fluoromicrograph of labelled axons in the RsC. Middle left, fluorescent images of vGluT1+ and PV+ target neurons in the retrosplenial cortex. Right, electronmicrographs of synaptic contacts established by BFPVNs with the targes cell types shown in the middle panels. Abbreviations: VDB, ventral limb of the diagonal band of Broca; MS, medial septum; RsG, retrosplenial cortex, granular part; Hipp, hippocampus; MMn, meidal mamillary nucleus; SUM, supramamillary nucleus; PT/sm, paratenial thalamic nucleus/stria medullaris; MO, medial orbital cortex; LSi, lateral septum, intermediate part; IL/DP, inflalimbic cortex/dorsal peduncular cortex; MMR, median raphe region; VTA, ventral tegmental area.

### HDB BFPVNs broadcast negative reinforcement signals to multiple downstream targets

Finally, we tested whether HDB BFPVNs send signals upon aversive US in a target-specific manner or distribute similar information to multiple downstream areas. To address this, we expressed GCaMP6s in PV-Cre mice (n = 12) by injecting AAV2/9.CAG.Flex.GCAMP6s.WPRE.SV40 bilaterally in the HDB and recorded their bulk calcium signals both at the cell body level within the HDB and in axonal projections in their main target areas such as the MS, the hippocampal CA1 and the retrosplenial cortex (Fig.6a). We recorded bulk calcium signals via fiber photometry while mice performed the Pavlovian conditioning task.

**Figure 6.**
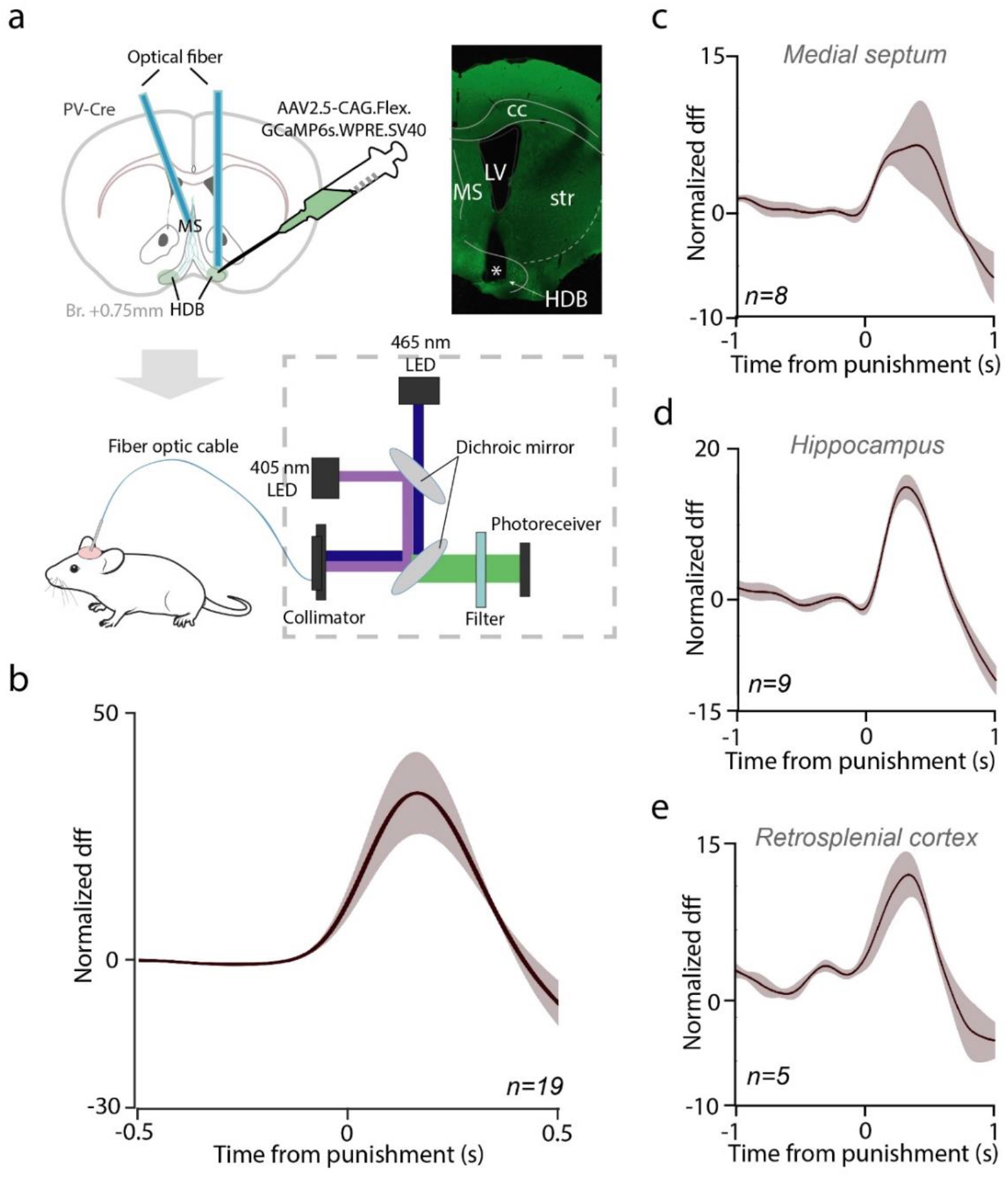
HDB BFPVNs broadcast negative feedback information to multiple limbic target regions. **a,** Schematic illustration of bulk calcium imaging of BFPVN activity in the HDB and in the main target areas. Top left, injection of GCaMP6s-expressing viral vector; top right, fluoromicrograph of an optical fiber track (green, GCaMP6s; asterisk, tip of the optical fiber). Bottom, schematic of bulk calcium imaging. **b,** Fiber photometry recordings of HDB BFPVN somata. Mean PETH of normalized dff aligned to punishment, recorded in the HDB (n = 19 sessions). **c,** Fiber photometry recordings of HDB BFPVN projections. Mean PETH of normalized dff aligned to punishment, recorded in the medial septum (n = 8 sessions). **d,** Mean PETH of normalized dff aligned to punishment, recorded in the hippocampus CA1 region (n = 9 sessions). **e,** Mean PETH of normalized dff aligned to punishment, recorded in the retrosplenial cortex (n = 5 sessions).

First, we confirmed that somatic calcium concentrations showed similar responses to air-puff punishments to what we observed when single neuron spiking responses of BFPVNs were analyzed (Fig.6b, Fig S6). Next, we found that BFPVN projections showed phasic punishment responses in all three target areas investigated (Fig.6c-e), arguing for a widespread broadcast- type function of HDB BFPVNs.

## Discussion

We demonstrated that PV-expressing long-range inhibitory neurons in the HDB nucleus of the basal forebrain respond strongly and homogeneously to punishments, broadcast this activity to multiple downstream targets in the limbic system, and that this activity aids the learning of stimulus-outcome associations. These results suggest that basal forebrain PV-expressing neurons are important for learning from negative experiences.

The BF contains a substantial GABAergic population that is approximately five times larger than its cholinergic population according to stereological estimates in rodents^4, 6^. This inhibitory population contains a significant long-range projecting neuron pool, many of which express parvalbumin^7, 8, 35–38^.

These projections include target areas implicated in cognitive processing, including frontal cortices and hippocampus^8, 10, 38^. In concert, non-specific lesions of the basal forebrain typically resulted in more substantial cognitive impairment compared to selective cholinergic lesions^39–,42^, suggesting a role for non-cholinergic BF neurons in associative learning^42, 43^. Also, the role of BF GABAergic neurons in eyeblink conditioning was demonstrated by a study applying selective GABAergic BF lesions^18^.

We therefore asked whether the PV-expressing population of BF GABAergic projection neurons, the BFPVNs, are essential for associative learning. We focused on the rostral portion of the horizontal nucleus of the diagonal band of Broca (HDB) and used bulk calcium imaging via fiber photometry and tetrode recordings combined with optogenetic tagging to measure BFPVN activity in the HDB. We first tested whether BFPVNs respond to behaviorally salient events during classical conditioning, while mice learned to associate previously neutral pure tone stimuli (CS) with water rewards and air puff punishments (US). Previous studies showed that a bursting population of BF non-cholinergic neurons respond to behaviorally relevant events irrespective of their valence during associative learning^19, 20^. We also found that BFPVNs responded to salient events including cue tones, reward and punishment; however, in contrast with Lin and colleagues, we demonstrated a strong preference of HDB BFPVNs towards responding to the aversive US, with only weaker and less consistent responses to CS and reward, suggesting that HDB BFPVNs constitute a specialized subpopulation of non-cholinergic BF neurons. We also found that BFPVNs markedly differed from cholinergic BF neurons, the BFCNs, in their responses: our previous studies revealed outcome prediction coding of BFCNs both following CS and US^21, 30, 31, 44^, absent in HDB BFPVNs.

The strong BFPVN responses to air-puff punishments suggest a role of these neurons in transmitting aversive information during associative learning. This is in line with a previous report suggesting that GABAergic medial septal-diagonal band neurons participate in nociceptive behaviors based on pharmacology and kainic acid induced lesions^45^. However, optogenetic activation of HDB BFPVNs did not elicit avoidance behavior in a conditioned place aversion test. In contrast, optogenetic inhibition of these neurons during air-puff punishments prevented mice from forming differential representation of CS cues based on their valence, demonstrated by a lack of differential anticipatory lick responses in optogenetically manipulated mice within the course of training. This lack of difference in CS-elicited licking was apparently due to an impaired inhibition of licking after Cue 2 that predicted likely punishment, in line with a previous study in which ibotenic acid injected to the BF of rats, preferentially destroying non-cholinergic neurons, lead to an increased false alarm rate in a visual detection task^14^.

What may be the input that conveys aversive information to HDB BFPVNs? To address this question, we mapped the input neurons to HDB BFPVNs by mono-transsynaptic retrograde tracing using rabies vectors. We found the largest density of input neurons in the lateral hypothalamus, known to transmit nociceptive information that it receives via direct monosynaptic connections from nociceptive neurons of the spinal chord and periaqueductal gray^46–48^. HDB BFPVNs receive inputs from different BF nuclei including the medial septum and the ventral pallidum, which may also transmit punishment-related signals^2, 21, 45, 49, 50^. Furthermore, the median raphe was also sending monosynaptic inputs to BFPVNs, raising the possibility of a direct brainstem source of aversive information^32, 51, 52^. This was supported by our finding that MRR input cells consisted mostly of vGluT2- and vGluT3-expressing glutamatergic neurons, known to play important roles in the brainstem control of negative experience^32, 34^. Importantly, the remaining major inputs to HDB BFPVNs were the lateral septum and the nucleus accumbens, thought to be important regulators of affect^53^. The potential convergence of aversive/affective information on BFPVNs revealed by rabies-based input mapping suggests that these neurons may act as an integrating and processing hub for aversive information.

We tested by anterograde tracing, fluorescence and electron microscopy to what target areas and neurons this aversive information is transmitted. HDB BFPVNs were projecting to neighboring BF nuclei, the vertical limb of the diagonal band of Broca and the medial septum (MS-VDB), where they synapsed on cholinergic and PV-expressing GABAergic neurons. They also sent dense projections to the hippocampus, where they formed synaptic contacts on PV- expressing and calretinin-expressing hippocampal interneurons. Of note, this innervation pattern was similar to what had been found for the septohippocampal GABAergic projection ^35, 54^. A third major target of HDB BFPVNs was the retrosplenial cortex, interfacing multimodal sensory and limbic information^55, 56^. Previous studies suggested that long-range projecting GABAergic BF neurons may be mostly disinhibitory^57–62^, which we confirmed in the CA1 and in the retrosplenial cortex, where we demonstrated HDB BFPVN synapses on PV-expressing interneurons; however, potential contacts onto pyramidal neurons could not be reliably tested. Further targets of HDB BFPVNs included other limbic structures like the mammillary and supramammillary nuclei and the paratenial thalamic nucleus, and to a lesser extent areas of the frontal cortex. This innervation pattern indicates a broad targeting of the limbic system by HDB BFPVNs, multiple areas of which were shown to be necessary for learning from aversive experience^63–70^.

Calcium imaging of HDB BFPVN projections performed at three major termination zones, the medial septum, the CA1 of the hippocampus and the retrosplenial cortex revealed that all HDB BFPVN projections exhibited similar response properties to what was observed at the cell body level. Specifically, all three projections showed prominent activation by air-puff punishments. Although some quantitative differences could be observed, it was not possible to dissect whether it was due to differences in axon density or differential response strength at the projection level. Altogether, the qualitatively similar response profiles suggest a broadcast function of HDB BFPVNs, sending similar message to broad areas of the limbic system.

BFPVNs have been shown to control cortical gamma oscillations^6, 9, 10^ and were suggested to participate in mediating arousal at different timescales, including brief and rapidly changing ‘microarousals’^6, 9, 71^. Relatedly, they were proposed to control changing levels of attention through cortical activation^6, 9, 21^. BFPVNs were also suggested to activate the default mode network, which is likely responsible for maintaining global oscillatory activity necessary for higher cognitive functions and conscious experience in humans, determining the significance of which will require future studies^72, 73^. Combining these and our results, we speculate that BFPVNs may rapidly activate limbic areas upon aversive stimuli to facilitate learning from negative experience. Thus, at least for aversive stimuli, BFPVNs might be the physical substrate of the ‘attention for learning’ concept^9, 74^.

## Methods

### Animals

Adult (over two months old) male Pvalb-IRES-Cre (PV-IRES-Cre or PV-Cre; n = 56, The Jackson Laboratory, RRID: IMSR_JAX:017320) mice were used for recording, optogenetic activation and inhibition, bulk calcium imaging and for immunohistochemistry, according to the regulations of the Hungarian Act of Animal Care and Experimentation (1998; XXVIII, section 243/1998, renewed in 40/2013) in accordance with the European Directive 86/609/CEE and modified according to the Directive 2010/63/EU. Experimental procedures were reviewed and approved by the Animal Welfare Committee of the Institute of Experimental Medicine, Budapest and by the Committee for Scientific Ethics of Animal Research of the National Food Chain Safety Office of Hungary (PE/EA/675-4/2016; PE/EA/1212-5/2017; PE/EA/864- 7/2019).

### Tetrode implantation surgery

Mice were implanted with microdrives housing 8 tetrodes and a 50 µm core optic fiber^21, 75, 76^. Mice were anesthetized with a ketamine-xylazine solution (83 mg/kg ketamine and 17 mg/kg xylazine, dissolved in 0.9% saline). The scalp was shaved and disinfected with Betadine, the skin and subcutaneous tissues were topically anesthetized with Lidocaine spray, and mice were placed in a stereotaxic frame (Kopf Instruments). Eyes were protected with eye ointment (Corneregel, Bausch & Lomb). The skin was opened with a single sagittal incision made by a surgical scalpel, the skull was cleaned, and a craniotomy was drilled above the horizontal limb of the diagonal band of Broca (HDB, antero-posterior 0.75 mm, lateral 0.60 mm; n = 4). Virus injection (AAV2/5.EF1a.Dio.hChR2(H134R)-eYFP.WPRE.hGH, Addgene, titer ≥ 1×10¹³ vg/mL; HDB, dorso-ventral 5.00 and 4.70 mm, 300 nl at each depth) and drive implantation was performed using the stereotaxic frame and a programmable nanoliter injector (Drummond Nanoject III). Ground and reference electrodes were implanted to the bilateral parietal cortex. The microdrive was secured in place using dental cement (Lang Dental). When the dental cement was cured, mice were transferred from the stereotaxic frame into their homecage and received analgesics (Buprenorphine, 0.1 mg/kg), local antibiotics (Gentamycin). Mice were closely monitored after surgery and were allowed 10-14 days of recovery before starting behavioral training.

### Anterograde tracing surgery

PV-IRES-Cre mice were injected with AAV2/8.CAG.Flex.eGFP (Addgene, titer ≥ 1×10¹³ vg/mL) anterograde tracing virus in the HDB (antero-posterior, 0.75 mm; lateral, 0.60 mm; dorso-ventral, 5.00 mm and 4.9 mm; 100 nl each side, injecting 50-50 nl each depth) using standard stereotactic surgery techniques described in the previous section. Following bilateral virus injection, craniotomies were closed with Vaseline, and scalp skin was sutured using standard surgical needle and thread (Dafilon). Postoperative care was applied as described above. The virus injection was followed by a 4-week virus expression period. Mice were then transcardially perfused (as described in details below) and their brains were processed for further immunohistology experiments.

### Retrograde tracing surgery

PV-IRES-Cre mice were anesthetized with 2% isoflurane followed by an intraperitoneal injection of a ketamine-xylazine anaesthetic mixture (83 mg/kg ketamine and 17 mg/kg xylazine, dissolved in 0.9% saline), then mounted in a stereotaxic frame (David Kopf Instruments). After exposing the skull surface, 20 nl of the Cre-dependent helper virus was injected to the HDB (bilateral, n = 2; unilateral, n = 1) using a Nanoliter 2010 microinjection pump (World Precision Instruments). The injection coordinates were defined by a stereotaxic atlas (Paxinos and Franklin, 2012) and were the following given in mm for the anteroposterior, mediolateral and dorsoventral axes, respectively: +0.75 (from Bregma), ±0.60 (from Bregma, unilateral in case of Mouse 3), -5.0 (from brain surface). After 3 weeks of survival, mice were injected with 20 nl of the G protein-deleted EnvA-pseudotyped rabies virus at the same coordinates. After 9 days of survival, mice were prepared for perfusion.

### Optical fiber implantation surgery for optogenetic activation, inhibition and fiber photometry recordings

PV-IRES-Cre mice were implanted using standard stereotactic surgery techniques described in the previous section. Following virus injection (for optogenetic activation: AAV2/5.EF1a.Dio.hChR2(H134R)-eYFP.WPRE.hGH, Addgene, titer ≥ 1×10¹³ vg/mL or AAV2/5.EF1a-eYFP.WPRE.hGH, Addgene, titer ≥ 7×10¹² vg/mL as control virus, 100 nl each side; for optogenetic inhibition: AAV2.9-CAG-Flex-ArchT-GFP, Addgene, titer ≥ 1×10¹³ vg/mL or AAV2.9-CAG-Flex-GFP, Addgene, titer ≥ 1×10¹³ vg/mL as control virus, 200 nl each side; for fiber photometry: AAV2/9.CAG.Flex.GCAMP6s.WPRE.SV40, Addgene, titer ≥ 1×10¹³ vg/mL, 300 nl each side; HDB, antero-posterior 0.75 mm, lateral 0.60 mm; dorso-ventral 5.00 and 4.7 mm). Following virus injection, optical fibers were implanted (for optogenetic activation: 200 µm core fiber was implanted unilaterally into HDB, antero-posterior 0.75 mm, lateral 0.60 mm; for optogenetic inhibition: 400 µm core fibers were implanted bilaterally in 0- and 20-degree angle into HDB; for fiber photometry: 400 µm core fibers were implanted into HDB and MS, antero-posterior 0.75 mm, lateral 1.2 mm, 20-degrees lateral angle; CA1 region of the hippocampus, antero-posterior -2.3 mm, lateral 1.4 mm, dorso-ventral 1.2 mm; retrospelnial cortex, antero-posterior -2.3 mm, lateral 0.5 mm, dorso-ventral 0.2 mm respectively). Mice received post-surgery care as described above in the *Tetrode implantation surgery* section and allowed 10-14 days of recovery before starting behavioral training.

### Probabilistic Pavlovian conditioning

PV-IRES-Cre mice were trained on an auditory Pavlovian conditioning task^22^ in a head-fixed behavioral setup that allowed millisecond precision of stimulus and reinforcement delivery^77^. On the first day of training, water restricted mice^22^ were head-fixed and given access to water reward whenever they licked a waterspout. Individual licks were detected by the animal’s tongue breaking an infrared photobeam. The following day, a 50 dB pure tone cue of one second duration was introduced that predicted likely reward (5 μl of water). After each cue presentation, water reward was delivered with 0.8 probability with a 200-400 ms delay, while the rest of the outcomes were omissions. The next trial started after the animal stopped licking for at least 1.5 s. The stimulus was preceded by a 1–4 s foreperiod according to a truncated exponential distribution, in order to prevent temporal expectation of stimulus delivery^78^. If the mouse licked in the foreperiod, the trial was restarted. We used the open source Bpod behavioral control system (Sanworks LLC, US) for operating the task.

Next, a second pure tone cue of well-separated pitch was introduced that predicted low probability reward (0.25). Next, air puff punishment (200 ms, 15 psi) was introduced in the following session with the final outcome contingencies (likely reward trials, 80% reward, 10% punishment, 10% omission; likely punishment trials, 25% reward, 65% punishment, 10% omission). These contingencies reflect careful calibration to keep mice motivated for the task. Different trial types (likely reward and likely punishment) and outcomes (water reward, air puff punishment and omission) were presented in a pseudorandomized order following the described outcome contingencies. Mice learned the task in approximately one week and consistently demonstrated reward anticipation by differential lick rate in response to the cues from the second week.

Auditory stimuli were calibrated using a precision electret condenser microphone (EMM-6, Daytonaudio) connected to a preamplifier digital converter (AudioBox iOne, PreSonus) and sound pressure levels were measured by the TrueRTA software^77^. Cue tone intensities were well above the hearing threshold for both sounds (20-30 dB for 4 and 12 kHz pure tones) but not as loud as to cause a startle reflex (around 70 dB)^21, 79^. Mice in this study underwent the same training protocol for consistency; however, we have reported that behavioral performance of the task did not depend on the identity (frequencies) of the conditioned stimuli (Figure S1 in ref. ^31^).

The aversive quality of air-puffs depends on the exact experimental settings. We applied 200 ms long puffs at 15 psi pressure (within the range of parameters used for eyeblink conditioning^80^). We demonstrated that mice consistently choose water without air-puff over water combined with air-puff, showing that air-puffs are aversive under these circumstances (see Figures 2C and 2D in ref. ^21^).

### Optogenetic inhibition of BFPVNs

A set of PV-IRES-Cre animals injected with AAV2.9-CAG-Flex-ArchT-GFP (Addgene, titer ≥ 1×10¹³ vg/mL, n = 6, ArchT group) or AAV2.9-CAG-Flex-GFP (Addgene, titer ≥ 1×10¹³ vg/mL, n = 8, Control group) were trained for a two-week period on the probabilistic Pavlovian conditioning task according to the training protocol described above^22^. In parallel with punishment presentation, a one-second-long yellow laser pulse (593 nm wavelength, Sanctity Laser) was delivered to the HDB bilaterally via optical fibers (400 µm core) to achieve optogenetic silencing of BFPVNs. Anticipatory lick rate (licking between cue onset and reinforcement delivery) was recorded in each (ArchT and Control) group.

### Conditioned place aversion task

PV-IRES-Cre animals injected with AAV2/5.EF1a.Dio.hChR2(H134R)-eYFP.WPRE.hGH (Addgene, titer ≥ 1×10¹³ vg/mL, n = 8, ChR group) or AAV2/5.EF1a-eYFP.WPRE.hGH (Addgene, titer ≥ 7×10¹^2^ vg/mL, n = 10, Control group) were placed in an arena with two sides, which were separated by a wall but connected with an open gate on the wall. The two sides of the arena were marked by different wall patterns (dotty or striped). In the arena, a center part (one third of the size of the arena) and a peripheral part was differentiated. The conditioning protocol took two days; on the first day, animals were placed into the chamber and were allowed to move freely for 5 minutes (habituation phase). The next day (test phase), mice were place into the chamber again and received 20 Hz laser light stimulation (473 nm wavelength, Sanctity Laser) on one of the sides. The stimulation side was assigned pseudorandomly across animals. Time spent in the center and peripheral part of the arena (in seconds or in % of total time spent in the arena), time spent on each side (in seconds or in % of total time spent in the arena), side crosses (number of crosses), velocity (cm/s) and different behavioral components (rearing, grooming, freezing, exploration) were measured.

### Tetrode recording

Extracellular recordings were performed with custom-built microdrives consisting of 8 movable nichrome tetrode electrodes (diameter, 12.7 μm; Sandvik) and a 50 μm core optical fiber (Thorlabs)^21, 75^. Microdrive screws were specifically designed to precisely control the descent of the tetrodes and the optic fiber into the brain; the pitch of the threading was 160 µm, therefore one eighths of a turn with the screw corresponded to 20 μm descent (M0.6 stainless steel flat head screw, 12 mm length; Easterntec, Shanghai, China). Before the tetrode surgery, we measured the protruding length of the electrodes on each microdrive with a micro-ruler (Olympus SZ61 stereomicroscope; micro-ruler, Electron Microscopy Tools). The electrodes were gold-plated (PEG was dissolved in DI water to create 1 g/l concentration, then 1.125 ml of PEG-solution was mixed with 0.375 ml gold plating solution, Neuralynx, see also Ferguson et al., 2009) to impedances of 30-100 kΩ measured at 1kHz (NanoZ) and dipped in DiI (ThermoFischer Scientific) red fluorescent dye to aid later track reconstruction efforts.

Before each recording session, the microdrive was connected (Omnetics) to a 32-channel RHD headstage (Intan). Data were digitized at 30 kHz and transferred from the headstage to a data acquisition board (Open Ephys) via a Serial Peripheral Interface cable (Intan). The tetrodes were advanced 0-100 µm after each recording session. The descent of the tetrodes was logged each recording day. Throughout the experiments, detailed notes of the assumed brain coordinates during each recording session were taken based on the length of the tetrodes measured before the surgery, stereotaxic information from the surgery and controlled screw turns on the microdrive.

### Optogenetic tagging

The optic fiber of the microdrive (see above) ended in an FC connector (Precision Fiber Products). The optic fiber was connected with an FC-APC patch chord during recording. Before the training session has begun, to optogenetically tag BFPVNs, 1 ms laser pulses were applied (473 nm, Sanctity) at 20 Hz for 2 seconds, followed by 3 seconds pause, repeated 20-30 times. Light-evoked spikes and incidental artifacts were monitored online using the OPETH plugin (SCR_018022)^82^ for Open Ephys and when light-induced photoelectric artifacts or population spikes were observed, laser power was adjusted as necessary to avoid masking of individual action potentials. Significance of photoactivation was assessed with offline data analysis by examining the SALT test based on spike latency distributions after light pulses, compared to a surrogate distribution using Jensen-Shannon divergence (information radius)^75, 83^. Neurons with p < 0.01 were considered light-activated, and thus BFPVNs. BFPVNs recorded on the same tetrode within 200 μm dorso-ventral distance were compared by waveform correlation and autocorrelogram similarity^49, 84^, and similar units were counted towards the sample size only once.

### Fiber photometry

A dual fiber photometry setup (Doric Neuroscience) was used to perform bilateral fluorescent calcium imaging. Imaging sessions were visualized using Doric Studio Software. Two LED light sources (465 nm, 405 nm) were amplitude-modulated by the command voltage of the two-channel LED driver (LEDD_2, Doric Neuroscience, the 465 nm wavelength light was modulated at 208 Hz and 405 nm wavelength was modulated at 572 Hz) and were channeled into fluorescent Mini Cubes (iFMC4, Doric Neuroscience). Light was delivered into the HDB, MS, CA1 or retrosplenial cortex via 400 µm core patch cord fibers that were connected to optical fiber implants during training sessions. The same optical fibers were used to collect the emitted fluorescence signal from the target areas. The emitted fluorescent signal was detected with 500–550 nm fluorescent detectors integrated in the Mini Cubes. Sampling rate of emitted signals was set to 12 kHz, decoded in silico and saved in a *.csv format.

### Perfusion

Mice were anesthetized with 2% isoflurane followed by an intraperitoneal injection of a mixture of ketamine-xylazine and promethazinium-chloride (83 mg/kg, 17 mg/kg and 8 mg/kg, respectively). After achieving deep anesthesia, mice were perfused transcardially (by placing the cannula into the ascending part of the aorta via an incision placed on the left ventricle wall) with saline for 2 minutes, followed by 4% paraformaldehyde (PFA) solution for 40 minutes, then saline for 10 minutes. After perfusion, mice were decapitated, and brains were carefully removed from the skull and postfixed in PFA overnight. Brains were then washed in PBS and cut into 50 μm thick coronal sections using a vibrating microtome (Leica VT1200S). Sections were thoroughly rinsed in 0.1 M PB (3 × 10 min) and used for further immunohistochemical experiments.

### Histological track reconstruction

After tetrode recording and fiber photometry experiments, animals were anesthetized with an intraperitoneal injection of ketamine-xylazine (83 mg/kg and 17 mg/kg, respectively) and underwent an electrolytic lesioning protocol (30 μA for 5s on two leads of two selected tetrodes, where opto-tagged BFPVNs were recorded; stimulator from Supertech, Pecs, Hungary). Twenty-to-thirty minutes after the electrolytic lesion, mice were perfused transcardially as described above. Microdrives and/or optical fibers were extracted and the brain was gently removed from the skull. The brain was postfixed in 4% PFA overnight and then washed in 0.1 M phosphate buffer (3 × 10 min). The explanted microdrives were examined under stereomicroscope and the protruding length of the electrodes were measured and compared to the depth registrations made during the recording sessions.

Coronal sections of 50 µm thickness were cut by a vibratome (Leica VT1200S). Special care was taken to section the brain perpendicular to brain surface, so that resulting sections could be easily aligned with coronal atlas images. The sections were thoroughly washed in phosphate buffer 3 times (10 min each) and mounted on microscopy slides in Aquamount mounting resin. Fluorescent micrographs of the sections were taken using a Nikon C2 confocal microscope. Four times four large field-of-view dark-field images, red and green fluorescent images were taken at 10x magnification in order to capture the whole sections.

Confocal microscopic images were further analyzed to reconstruct in vivo recording locations. The recording locations were referenced to atlas coordinates^85^. By referencing recording location to atlas coordinates, individual size differences of mouse brains compared to the atlas reference and slight deviations from the vertical direction during electrode descent were accounted for. Dark-field whole-section brain images aided atlas alignment, since they provided a detailed contrast for white and grey matter. By using Euclidean transformations, atlas images of coronal sections were morphed on the corresponding dark-field brain images, to verify area boundaries and determine the coronal plane of the section. If the brain section was non-uniformly distorted during histological processing, special care was taken to accurately map the vicinity of the electrode tracks within the target areas. Next, green fluorescent images of the same sections were transformed similarly as the dark-field images to verify PV expression within the boundaries of anatomical structures corresponding to the atlas images. In case of tetrode recordings, aligned atlas images were superimposed on red fluorescent images of the same field-of-view, which showed the DiI-labeled electrode tracks. Coordinates of electrode entry points and deepest points in the brain were marked by DiI tracks and the tip of the electrode was localized using small electrolytic lesions. Coordinates of electrode endpoints were read from the atlas image and were used to interpolate the recording locations referenced to the atlas coordinates, based on experimental logs of the protrusion of the electrode. Based on this interpolation procedure, antero- posterior, lateral and dorso-ventral coordinates and an atlas-defined brain area were assigned to each recording session and thus to each recorded neuron.

Sections from mice implanted with optic fibers were processed similarly; dark-field whole section brain images were used for atlas alignment, while green fluorescent images verified the expression of ChR2, ArchT or GCaMP6s. Optic fiber locations were reconstructed based on the tissue track caused by the fiber.

### Anterograde tracing

Sections were washed in PB (0.1 M) and immersed in 30% sucrose solution overnight, then freeze-thawed over liquid nitrogen three times. Sections were extensively washed in 0.1M PB and TBS and blocked in 1% human serum albumin (HSA; Sigma-Aldrich) solution for 1 h. Then, sections were incubated in primary antibodies against choline-acetyltransferase (ChAT) or parvalbumin (PV) or calretinin (CR) together with anti-EGFP antibody (see Table S2) for 48-60 hours. Sections were rinsed 3 times for 10 minutes in TBS; secondary fluorescent antibodies (see Table S3) were applied overnight. Sections were rinsed in TBS and 0.1 M PB and mounted on slides in Aquamount mounting medium (BDH Chemicals Ltd). Next, fluorescent images were taken with a Nikon A1R Confocal Laser Scanning Microscope. In target areas of BFPVNs, fluorescent images of 10x magnification were taken. The axon densities of BFPVN projections were estimated by the mean pixel brightness value in each region.

### Retrograde tracing

Sections were washed in 0.1 M PB, incubated in 30% sucrose overnight for cryoprotection, then freeze-thawed over liquid nitrogen three times for antigen retrieval. Sections were subsequently washed in PB and Tris-buffered saline (TBS) and blocked in 1% HSA in TBS, then incubated in a mixture of primary antibodies for 48-72 hours. This was followed by extensive washes in TBS, and incubation in the mixture of appropriate secondary antibodies overnight. Then, sections were washed in TBS and PB, dried on slides and covered with Aquamount.

The combinations and specifications of the primary and secondary antibodies used are listed in TableS2 and TableS3. The specificities of the primary antibodies were extensively tested, using knock-out mice if possible. Secondary antibodies were tested for possible cross-reactivity with the other antibodies used, and possible background labeling without primary antibodies was excluded.

Sections were imaged using a Nikon Ni-E Eclipse epifluorescent microscope system equipped with a 10× air objective and a Nikon DS-Fi3 camera. Fifty μm thick coronal sections spaced at 300 μm were prepared, so that approximately one-sixth of the whole brain was sampled to measure and estimate the number of transsynaptically labeled input cells.

### Correlated light an electron microscopic analysis

Sections from the anterograde tracing experiments (see above) were imaged with Nikon A1R confocal laser scanning microscope for axon density measurements and for imaging target cell types. First, sections from anterograde tracing experiments were used for immunohistochemical staining against markers of possible target cell types (nNOS, neuronal nitrogen-monoxid synthase; PV; VIP, vasoactive intestinal peptide; SOM, somatostatin; CB, calbindin; CR, calretinin; ChAT). A subset of sections were mounted in Aquamount mounting medium and imaged at 60x magnification and oil immersion with A1R Nikon Confocal laser scanning microscope. Sections from at least 3 animals were examined for possible contact sites between BFPVN projecting axons (labelled by eGFP) and target cell types. When possible terminal contacts were observed with a target cell type, another subset of sections from the same immunohistochemical experiment were mounted on slides in 0.1 M PB and were imaged similarly as described above. Possible terminal contacts were photographed at multiple magnifications (60x, 20x, 10x, and 4x) and surrounding landmarks blood vessels, location of ventricles, corpus callosum and other white matter tracts, tears at the edge of the section) were marked. After imaging, the sections were de-mounted from the slides to perform a second immunohistochemical staining to label eGFP (labeling BFPVNs and their projections) with 3,3’-Diaminobenzidine (DAB) as chromogen. Sections were rinsed extensively in TBS (3 × 10 minutes), then anti-chicken biotinylated secondary antibody was applied (in order to label eGFP-containing fibers, which were previously recognized by a chicken anti-eGFP primary antibody, see above) overnight. This step was followed by incubation in avidin–biotinylated horseradish peroxidase complex overnight (Elite ABC; 1:500; Vector Laboratories); the immunoperoxidase reaction was developed using DAB (Sigma-Aldrich) chromogen. Next, sections were mounted on slides in PB and light microscopic imaging (Nikon C2) was performed to reveal BFPVN axons. Next, light microscopic images were taken and aligned with former fluorescent images based on previously marked landmarks to determine the precise location of possible contact sites between BFPVN axons and target cell types on the light microscopic images. After this, sections were incubated with 0.5 % osmium tetroxide in 0.1 M PB on ice and they were dehydrated using ascending alcohol series and acetonitrile, finally incubated in 1% uranylacetate in 70% ethanol and embedded in Durcupan resin (ACM; Fluka). Brain regions containing the possible contact sites were re-embedded into resin blocks and 70 nm thick sections were cut with an ultra-microtome (Leica EM UC 6). Electron microscopic images were taken either with a Hitachi H-7100 transmission electron microscope or a FEI series electron microscope to reveal synaptic contacts between BFPVNs and target cell types in the MS, CA1 and retrosplenial cortex.

### Data analysis

#### Analysis of extracellular recordings

Data analysis was carried out using custom written Matlab code (R2016a, Mathworks). Tetrode recording channels were digitally referenced to a common average reference, filtered between 700-7000 Hz with Butterworth zero-phase filter and spikes were detected using a 750 ms censoring period. Action potentials were sorted into putative single neurons manually by using MClust 3.5 (A.D Redish, University of Minnesota). Autocorrelations were examined for refractory period violations and only putative units with sufficient refractory period were included in the data set. Neurons with p < 0.01 significance values of the SALT statistical test (H-index) were considered light-activated, therefore BFPVNs (see above). Spike shape correlations between light induces and spontaneous spikes were calculated for all BFPVNs. Correlation coefficient exceeded R = 0.84 in all and R = 0.9 in 22/36 optotagged neurons (0.96 ± 0.0085, median ± SE; range, 0.8427 – 0.9989, see also Cohen et al., 2012). Only those BFPVNs with high cluster quality (isolation distance > 20 and L-ratio < 0.15 calculated based on full spike amplitude and first principal component of the waveform) were included in the final dataset for further analysis^21, 87^.

After spike sorting, the activity of individual neurons was aligned to cue presentation, reward and punishment delivery. Statistics were carried out on each neuronal unit; baseline activity was defined by taking a 1 s window before the cue, then firing rate in the baseline window was compared to firing rate in the test window (0-0.5 s after the event). One-sided hypotheses of firing rate increase and decrease were tested by Mann-Whitney U-test (p < 0.001; for cue-evoked activity, separately for likely reward and likely punishment cue). Neurons were sorted into different groups based on their statistically significant responses to the behaviorally relevant events (e.g. activated by cue, inhibited by reward etc.).

We calculated event-aligned raster plots and peri-event time histograms (PETHs) for all neurons, aligned to both behavioral events and photostimulation pulses. To calculate average PETHs, individual PETHs were Z-scored by the mean and standard deviation of a baseline window (1s before cue onset) and averaged across neurons. Response latency and jitter to optogenetic stimulation were determined based on activation peaks in the PETHs.

Autocorrelograms (ACG) were calculated at 0.5 ms resolution. Burst index (BI) was calculated by the normalized difference between maximum ACG for lags 0-10 ms and mean ACG for lags 180-200 ms, where the normalizing factor was the greater of the two numbers, yielding an index between -1 and 1^88^. A neuron with a BI > 0.3 and refractory period <2 ms was considered bursting, confirmed by the presence of ‘burst shoulders’ on average ACG in the ‘bursting group’ and the complete lack of ‘burst shoulders’ on the average ACG in the ‘non-bursting’ group. To examine burst coding in BFPVNs, analysis of neuronal responses to punishment were also carried out when only burst spikes or single spikes were considered for a neuron. A burst was detected whenever an inter-spike interval (ISI) was < 10 ms and subsequent spikes were considered as part of the burst as long as the ISI remained < 15 ms. Data were Z-score normalized with their surrogate mean and standard deviation to plot average ACG. The surrogates were generated according to^89^.

#### Fiber photometry signal processing

Fiber photometry signals were preprocessed according to^90^. Briefly, the recorded fluorescence signals were filtered using a low-pass Butterworth digital filter below 20 Hz to remove high frequency noise. To calculate dff, a least-squares linear fit was applied to the isosbestic 405 nm signal to align its baseline intensity to that of the calcium-dependent 465 nm (f465) signal. The fitted 405 nm signal (f_405,fitted_) was used to normalize the 465 nm signal as follows:

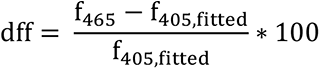

to remove the effect of motion and autofluorescence. A 0.2 Hz high pass Butterworth digital filter was used to filter out the slow decrease of the baseline activity observed during the recording session. Finally, the dff signal was aligned to cue and reinforcement times, Z-scored by the mean and standard deviation of a baseline window (1s before cue onset) and averaged across sessions.

### Statistics

Firing rates and other variables were compared across conditions using non-parametric tests (normality of the underlying distributions could not be determined unequivocally). For non-paired samples, Mann-Whitney U-test was used; for paired samples, Wilcoxon signed-rank test was applied (two-sided unless stated otherwise). Pearson’s correlation coefficient was used to estimate correlations between variables, and a standard linear regression approach was used to judge their significance (one-sided F-test, in accordance with the asymmetric null hypothesis of linear regression). Peri-event time histograms (PETHs) show mean ± SE. Box-whisker plots show median, interquartile range and non-outlier range, with all data point overlaid. Bar graphs show mean of paired variables and all data points overlaid.

ROC analysis was performed to test if the anticipatory lick rate difference between Cue 1 and Cue 2 trials was different between control and ArchT groups in the optogenetic inhibition experiment. First, average PETH of lick rates were calculated for control and ArchT groups in a 1.1 s long time window from cue onset (from cue to reinforcement), then values at every 10 ms of the time window were taken. Next, area under the ROC curve (auROC) based on the cumulative density functions of lick rate differences in control and ArchT groups were calculated for each time point. The significance of the auROC was tested with a bootstrap permutation test with 200 resamplings.

## Data availability

Data are available from the corresponding author upon reasonable request.

## Code availability

MATLAB codes generated for this study are available at https://github.com/hangyabalazs/PV_Pavlovian_analysis

## Acknowledgements

We thank Katalin Lengyel for technical assistance in anatomical methods, Ilka Jakab and Melinda Szabó for assistance in behavioral training and Dr. Norbert Hájos for helpful comments on the manuscript. We thank the FENS-Kavli Network of Excellence for fruitful discussions. This work was supported by the “Lendület” Program (LP2015-2/2015), NKFIH K135561, and the European Research Council Starting Grant no. 715043 to BH; the Hungarian Brain Research Program NAP3.0 (NAP2022-I-1/2022) by the Hungarian Academy of Sciences and the European Union project RRF-2.3.1-21-2022-00004 within the framework of the Artificial Intelligence National Laboratory to BH and GN; Frontline Research Excellence Programme by the Hungarian National Research, Development and Innovation Office (NRDI Fund 133837) and the European Union project RRF-2.3.1-21-2022-00011 within the framework of the Translational Neuroscience National Laboratory to GN; ÚNKP-21-3 New National Excellence Program of the Ministry for Innovation and Technology and the Kerpel Scholarship of Semmelweis University to P.H. We acknowledge the help of the Nikon Center of Excellence at the Institute of Experimental Medicine (IEM), Nikon Europe, Nikon Austria, and Auro-Science Consulting for kindly providing microscopy support and the supportive help of the Central Virus Laboratory and the Behavioral Unit of IEM. We thank Mackenzie Mathis for open access science art at SciDraw.

## Author contributions

P.H. and B.H. conceived the study; P.H. performed the immunohistological experiments, fluorescent and electron microscopic imaging; M.M and G.N. performed the retrograde tracing experiments; P.H. and Z.Z. performed the recording and optogenetic activation experiments; P.H., A.V. and V.L. conducted the fiber photometry and optogenetic inhibition experiments; P.H. and V.L. analyzed the data; P.H., M.M. and G.N. prepared the figures; B.H. and P.H. wrote the manuscript with inputs from all authors.

## Supplementary information

**Figure S1.**
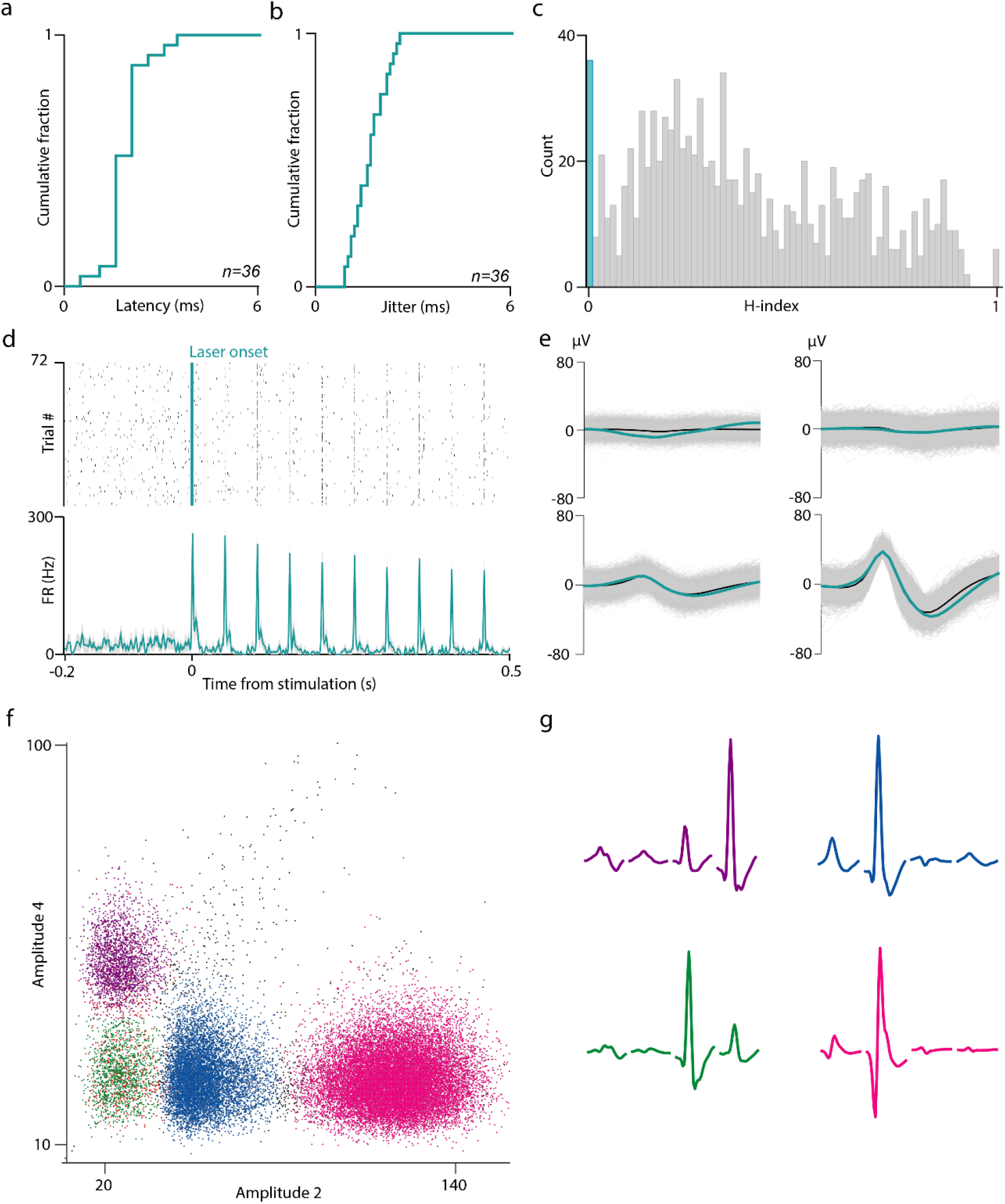
Optogenetic tagging of HDB BFPVNs. **a,** Cumulative histogram of the peak response latency of BFPVNs after optogenetic stimulation (n = 36). **b,** Cumulative histogram of the jitter of BFPVN spike responses after optogenetic stimulation (n = 36). **c,** Distribution of the significance values of the SALT statistical test (H-index) for all recorded neurons (blue, p < 0.01, tagged BFPVNs; grey, p > 0.01, untagged neurons). **d,** Example spike raster and PETH of an optogenetically tagged BFPVN responding to 20 Hz blue laser light stimulation. **e,** Right, average spike waveform of the same BFPVN on the four tetrode channels (blue, average light- evoked spikes; black, average spontaneous spikes; grey, all spikes). **f,** Spike clusters plotted in feature space from an example recording session. **g,** Average spike waveforms of the recorded neurons on each tetrode channel from the same session.

**Figure S2.**
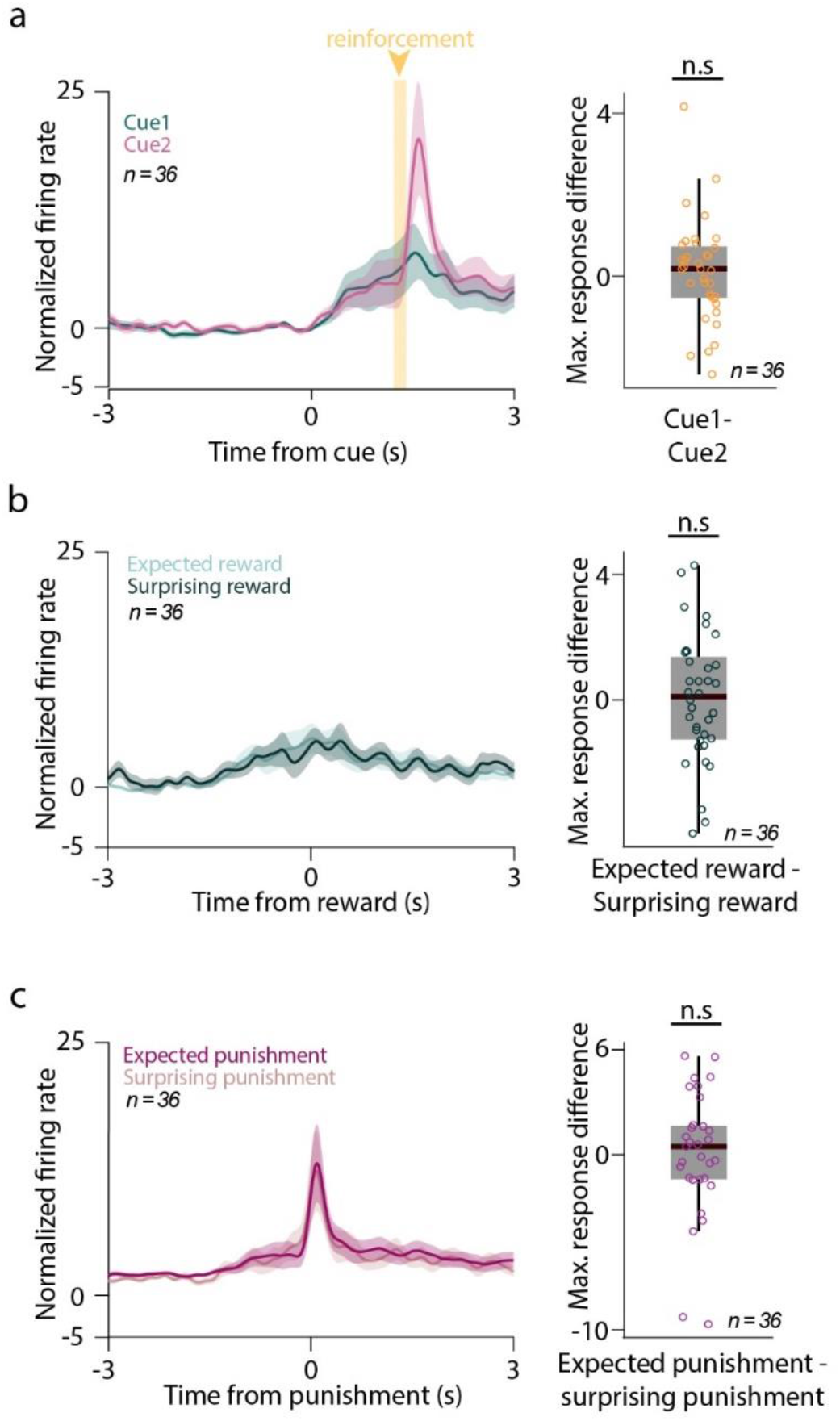
HDB BFPVNs are not modulated by outcome expectation. **a,** Left, average PETH of BFPVNs aligned to cue onset (n = 36). Right, difference of peak response to Cue 1 and Cue 2 (in a 0-0.5s time window from cue onset). n.s., p > 0.05, p = 0.6151, Wilcoxon signed-rank test. **b,** Left, average PETH of BFPVNs aligned to expected and surprising reward (n = 36). Right, difference of peak response to expected and surprising reward. n.s., p > 0.05, p = 0.8628, Wilcoxon signed-rank test. **c,** Left, average PETH of BFPVNs aligned to expected and surprising punishment (n = 36). Right, difference of peak response to expected and surprising punishment. n.s., p > 0.05, p = 0.5566, Wilcoxon signed-rank test.

**Figure S3.**
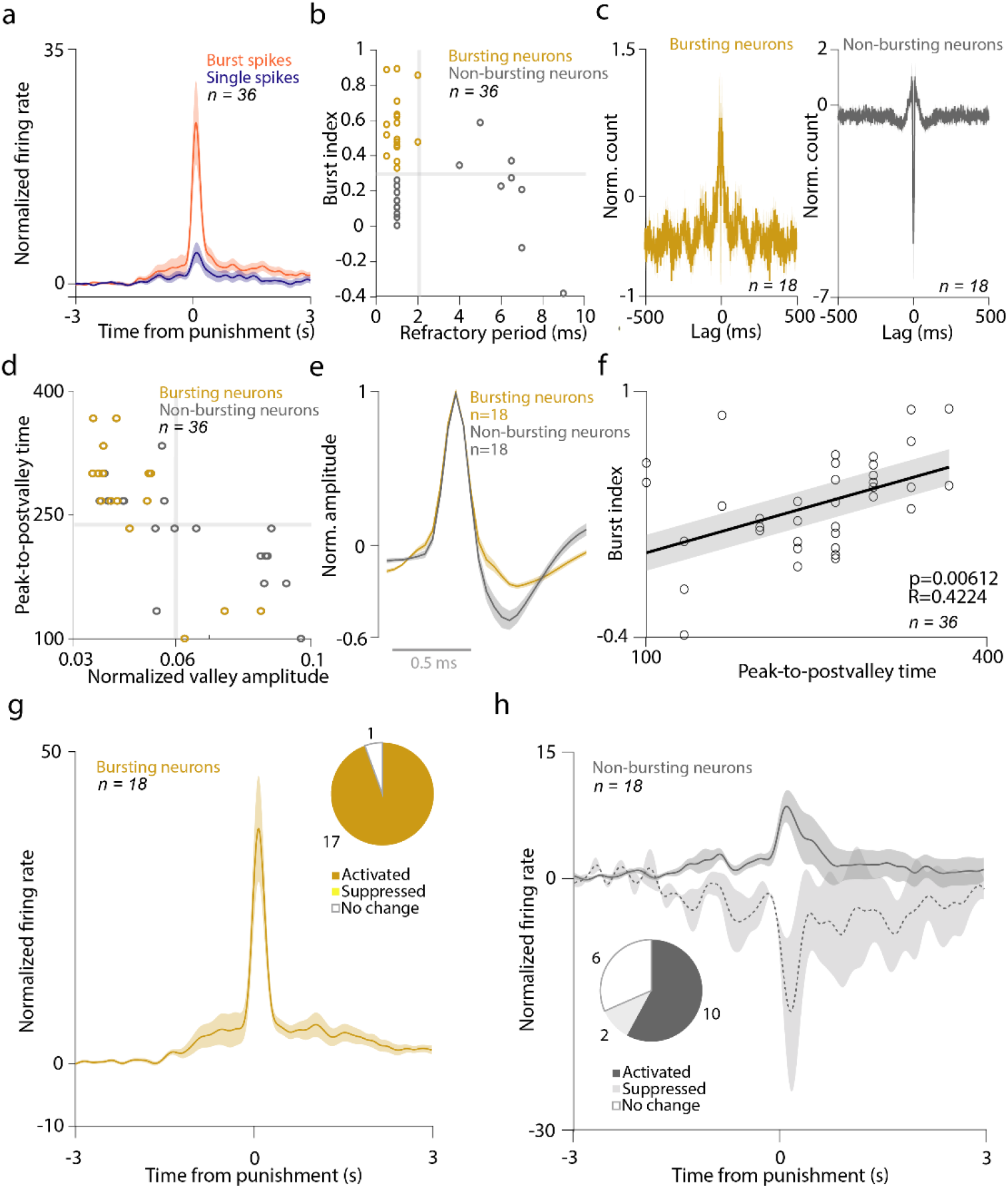
Electrophysiological properties of HDB BFPVNs. **a,** Average PETH of burst spikes and single spikes aligned to punishment onset (n = 36). **b,** BFPVNs were partitioned based on burst index and refractory period. Neurons with high (> 0.3) burst index and short (< 2 ms) refractory period were considered as bursting neurons. Neurons with low burst index and/or long refractory period were considered non-bursting. **c,** Average autocorrelogram of bursting (left) and non-bursting (right) neurons. **d,** Spike shape features of BFPVNs (peak-to- post-valley time and post-valley magnitude, normalized to the integral). Most bursting neurons had smaller valley and longer peak-to-post-valley time. **e,** Average spike shape of bursting and non-bursting neurons. **f,** Correlation of burst index and peak-to-post-valley time. **g,** Average PETH of bursting BFPVNs aligned to punishment. Pie chart showing activation, suppression or no response to punishment of bursting BFPVNs (n = 18). **h,** Average PETH of non-bursting BFPVNs aligned to punishment. Pie chart showing activation, suppression or no response to punishment of non-bursting BFPVNs (n = 18).

**Fig S4.**
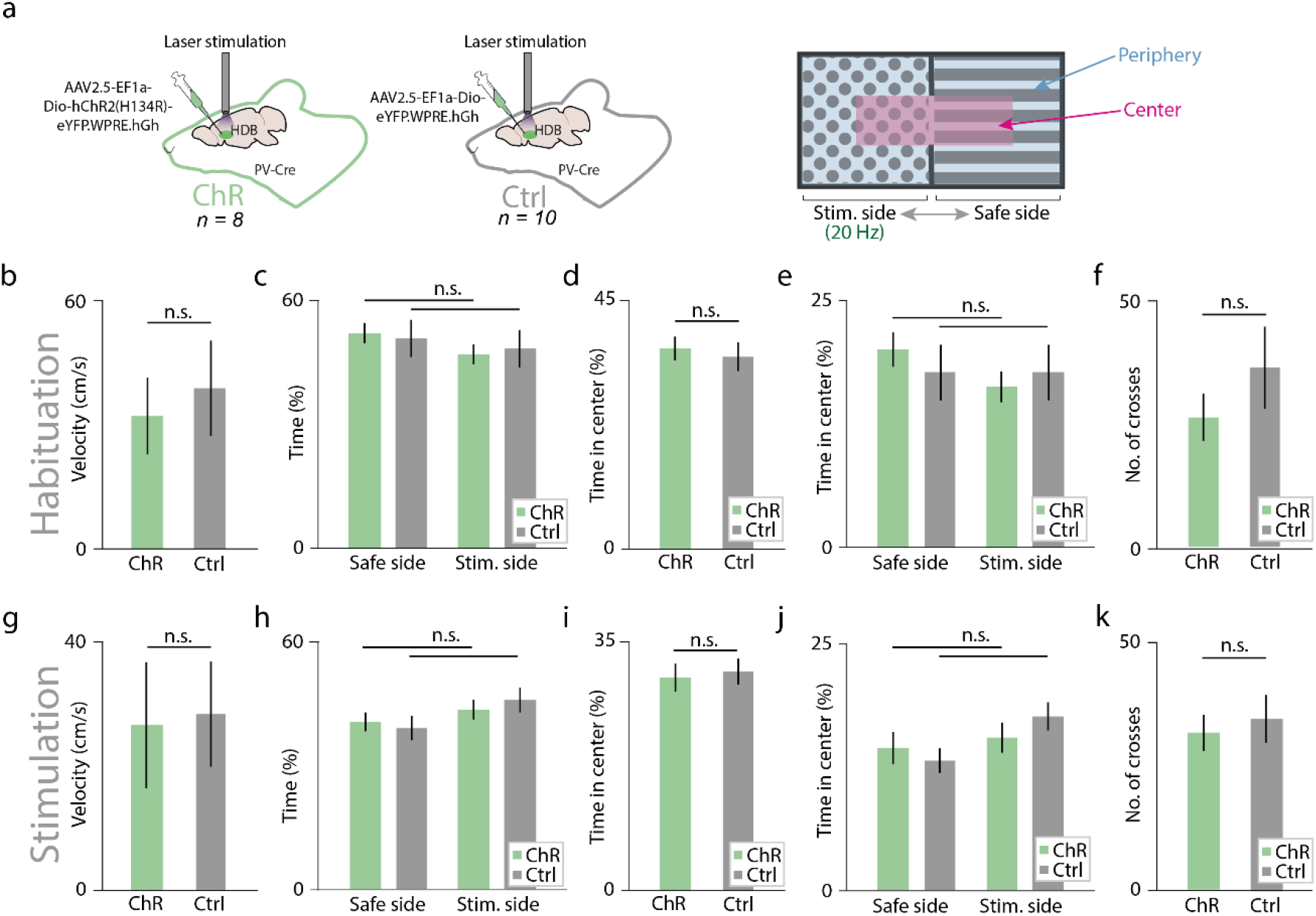
Conditioned place aversion. **a,** Schematic illustration of the experiment (for further details, see Methods). ChR, channelrhodopsin; Ctrl, control; Stim. side, stimulated side. **b,** Velocity of channelrhodopsin-expressing (ChR, n = 8) and control (Ctrl, n = 10) animals during habituation (n.s., p > 0.05, p = 0.4557, Mann-Whitney U-test). All bar graphs in this figure show mean ± SEM **c,** Proportion of time spent on the safe (non-stimulated) and stimulated side during habituation (n.s., p > 0.05, p = 0.9591 and p = 0.6454, Mann-Whitney U-test). **d,** Overall proportion of time spent in the center area during habituation (n.s., p > 0.05, p = 0.8673, Mann-Whitney U-test). **e,** Proportion of time spent in center area on the stimulated and safe sides during habituation (n.s., p > 0.05, p = 0.2134 and p = 0.7643, Mann-Whitney U-test). **f,** Number of side crosses during habituation (n.s., p > 0.05, p = 0.4538, Mann-Whitney U-test). **g-k,** Same as in B-F, but during the stimulation phase (n.s., p > 0.05, p = 0.6334, p = 0.7985, p = 0.995, p = 0.6734, p = 0.4352, p = 0.2452, p = 0.3955, respectively, Mann-Whitney U-test).

**Fig S5.**
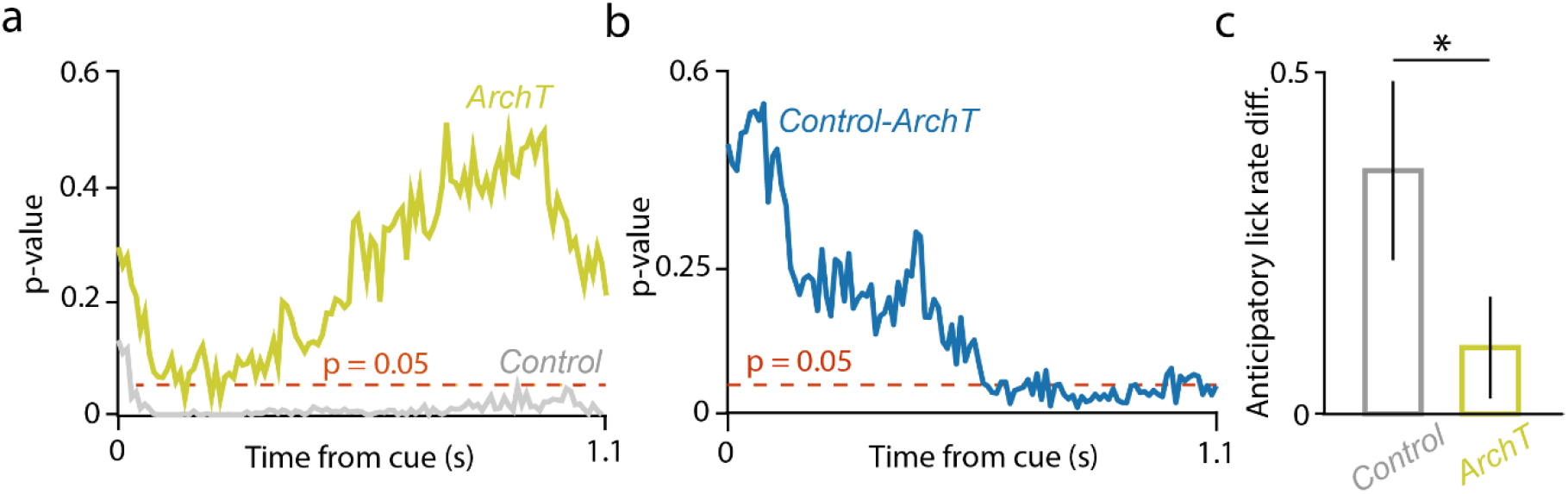
ROC analysis of anticipatory lick rate difference in control and ArchT groups. **a,** Statistical significance of cue-specific anticipatory lick rate difference as a function of time from cue onset (ROC analysis, see Methods). **b,** Statistical significance of the cue-related anticipatory lick rate difference between the ArchT and the control group as a function of time from cue onset (ROC analysis, see Methods). **c,** Bar graph showing mean ± SEM of anticipatory lick rate difference in control (left) and ArchT (right) groups in a 0.6-1.1 s window from cue onset. *, p < 0.05, p = 0.03669, Mann-Whitney U-test.

**Fig S6.**
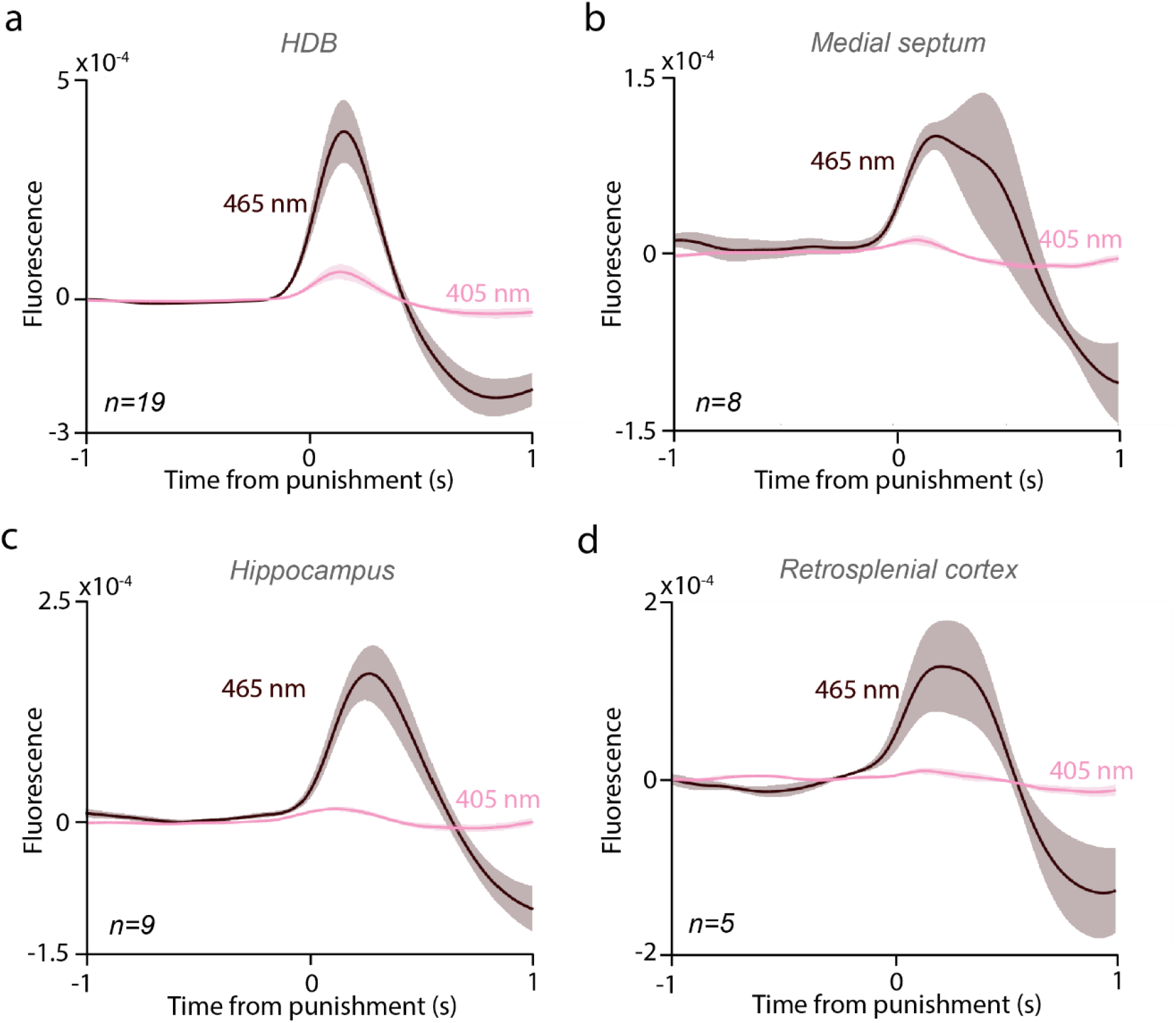
Isosbestic channels of fiber photometry recordings. **a,** PETH of 465 nm and 405 nm wavelength fluorescent signals aligned to punishment, recorded in the HDB (n = 19 sessions). **b,** PETH of 465 nm and 405 nm wavelength fluorescent signals aligned to punishment, recorded in the MS (n = 8 sessions). **c,** PETH of 465 nm and 405 nm wavelength fluorescent signals aligned to punishment, recorded in the CA1 hippocampus (n = 9 sessions). **d,** PETH of 465 nm and 405 nm wavelength fluorescent signals aligned to punishment, recorded in the RsC (n = 5 sessions).

**Table S1.**
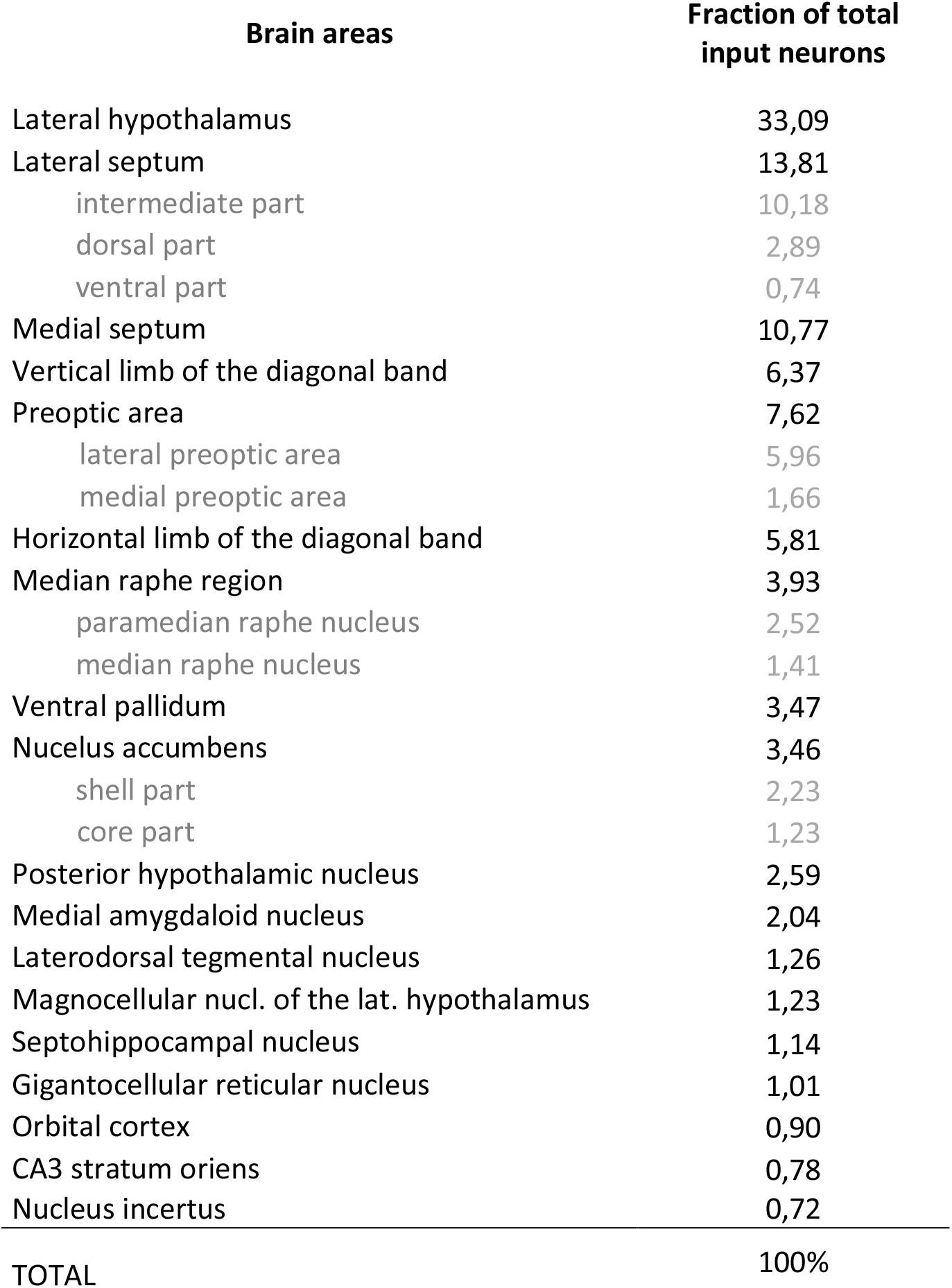
Fraction of total inputs to HDB BFPVNs.

| Brain areas | Fraction of total<br>input neurons |
| --- | --- |
| Lateral hypothalamus | 33,09 |
| Lateral septum | 13,81 |
| intermediate part | 10,18 |
| dorsal part | 2,89 |
| ventral part | 0,74 |
| Medial septum | 10,77 |
| Vertical limb of the diagonal band | 6,37 |
| Preoptic area | 7,62 |
| lateral preoptic area | 5,96 |
| medial preoptic area | 1,66 |
| Horizontal limb of the diagonal band | 5,81 |
| Median raphe region | 3,93 |
| paramedian raphe nucleus | 2,52 |
| median raphe nucleus | 1,41 |
| Ventral pallidum | 3,47 |
| Nucleus accumbens | 3,46 |
| shell part | 2,23 |
| core part | 1,23 |
| Posterior hypothalamic nucleus | 2,59 |
| Medial amygdaloid nucleus | 2,04 |
| Laterodorsal tegmental nucleus | 1,26 |
| Magnocellular nucl. of the lat. hypothalamus | 1,23 |
| Septohippocampal nucleus | 1,14 |
| Gigantocellular reticular nucleus | 1,01 |
| Orbital cortex | 0,90 |
| CA3 stratum oriens | 0,78 |
| Nucleus incertus | 0,72 |
| TOTAL | 100% |

**Table S2.** Primary antibodies used in immunhistochemical experiments.

| Antigen | Host | Dilution | Source | Catalog number | Specificity | Characterized in |
| --- | --- | --- | --- | --- | --- | --- |
| eGFP | Chicken | 1:2000 | Thermo Fisher Scientific | A10262 | No staining in mice not injected with eGFP-expressing virus. | Information of the distributor. |
| mCherry | Rabbit | 1:2000 | BioVision | 5993-100 | No staining in mice not injected with mCherry-expressing virus. | Information of the distributor. |
| vGluT3 | Guinea pig | 1:1000 | Frontier Institute | 522588 | Immunoblot detects a single protein band at 60-62 kDa. | Information of the distributor. |
| RFP | Rat | 1:2000 | Chromotek | 5F8 | No staining in mice not injected with mCherry-expressing virus. | Information of the distributor. |
| 5-HT | Rabbit | 1:10000 | Immunostar | 20080 | KO verified. | <sup>91</sup> |
| Chat | rabbit | 1:1000 | Synaptic Systems | 297013 | Specific for rat and mouse Chat. | Information of the distributor. |
| Chat | goat | 1:1000 | Merck | AB-144P | Specific for Chat in mouse among other species. | Information of the distributor. |
| PV | rabbit | 1:1000 | Swant | PV27 | KO verified. | Information of the distributor. |
| CR | rabbit | 1:10000 | Swant | 7697 | KO verified. | Information of the distributor. |

**Table S3.**
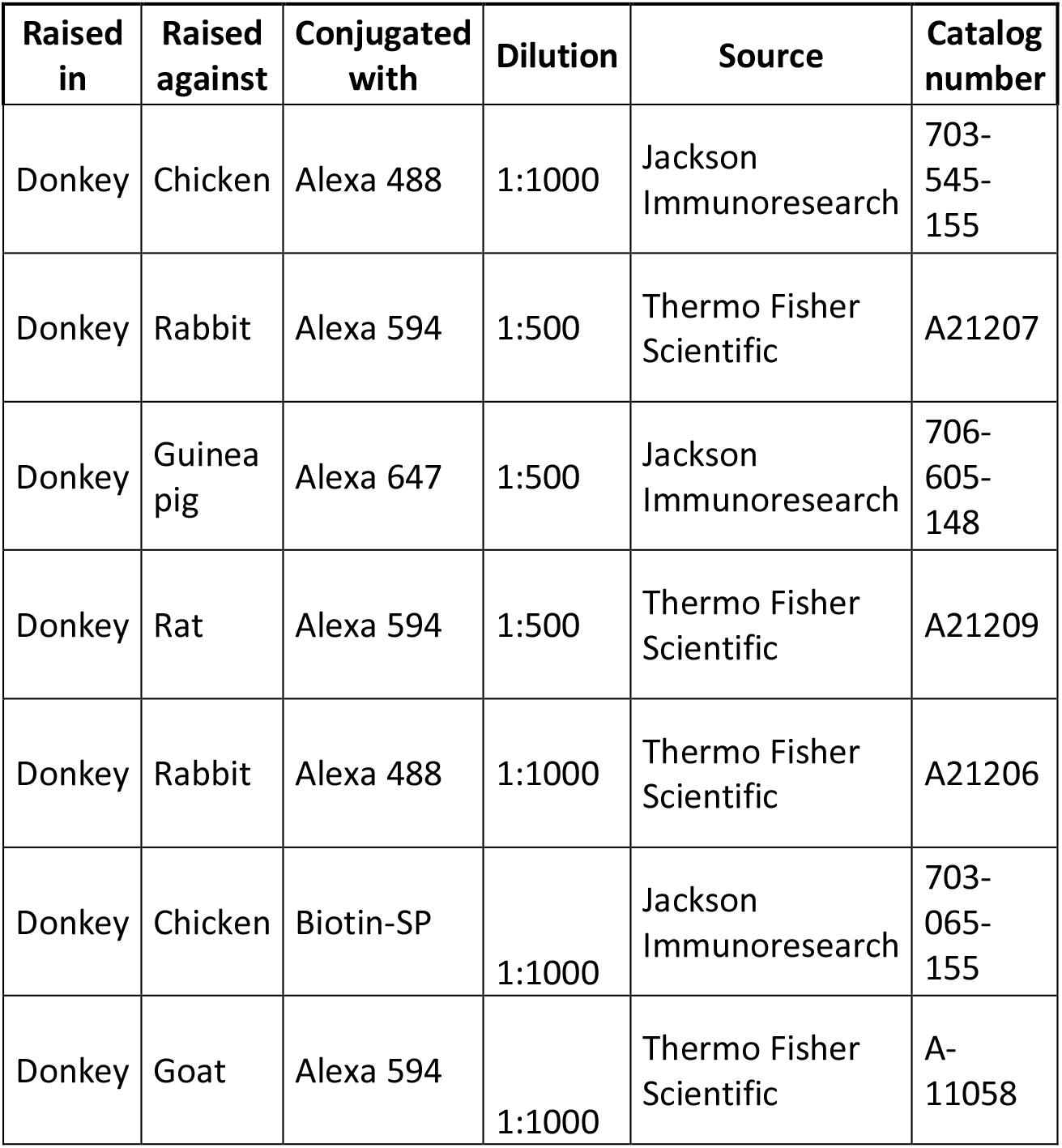
Secondary antibodies used in immunohistochemical experiments.

| **Raised in** | **Raised against** | **Conjugated with** | **Dilution** | **Source** | **Catalog number** |
| --- | --- | --- | --- | --- | --- |
| Donkey | Chicken | Alexa 488 | 1:1000 | Jackson ImmunoResearch | 703-545-155 |
| Donkey | Rabbit | Alexa 594 | 1:500 | Thermo Fisher Scientific | A21207 |
| Donkey | Guinea pig | Alexa 647 | 1:500 | Jackson ImmunoResearch | 706-605-148 |
| Donkey | Rat | Alexa 594 | 1:500 | Thermo Fisher Scientific | A21209 |
| Donkey | Rabbit | Alexa 488 | 1:1000 | Thermo Fisher Scientific | A21206 |
| Donkey | Chicken | Biotin-SP | 1:1000 | Jackson ImmunoResearch | 703-065-155 |
| Donkey | Goat | Alexa 594 | 1:1000 | Thermo Fisher Scientific | A-11058 |

## References

1. Mesulam, M.-M., Mufson, E. J., Levey, A. I. & Wainer, B. H. Cholinergic innervation of cortex by the basal forebrain: Cytochemistry and cortical connections of the septal area, diagonal band nuclei, nucleus basalis (Substantia innominata), and hypothalamus in the rhesus monkey. J. Comp. Neurol. 214, 170–197 (1983).

2. Zaborszky, L., van den Pol, A. & Gyengesi, E. The Basal Forebrain Cholinergic Projection System in Mice. in The Mouse Nervous System (eds. Watson, C., Paxinos, G. & Puelles, L.) 684–718 (Elsevier, 2012). doi:10.1016/B978-0-12-369497-3.10028-7.

3. Lehmann, J., Nagy, J. I., Atmadja, S. & Fibiger, H. C. The nucleus basalis magnocellularis: The origin of a cholinergic projection to the neocortex of the rat. Neuroscience 5, 1161–1174 (1980).

4. Gritti, I. et al. Stereological estimates of the basal forebrain cell population in the rat, including neurons containing choline acetyltransferase, glutamic acid decarboxylase or phosphate-activated glutaminase and colocalizing vesicular glutamate transporters. Neuroscience 143, 1051–64 (2006).

5. Gritti, I., Manns, I. D., Mainville, L. & Jones, B. E. Parvalbumin, calbindin, or calretinin in cortically projecting and GABAergic, cholinergic, or glutamatergic basal forebrain neurons of the rat. J. Comp. Neurol. 458, 11–31 (2003).

6. Yang, C., Thankachan, S., McCarley, R. W. & Brown, R. E. The menagerie of the basal forebrain: how many (neural) species are there, what do they look like, how do they behave and who talks to whom? Curr. Opin. Neurobiol. 44, 159–166 (2017).

7. Agostinelli, L. J., Geerling, J. C. & Scammell, T. E. Basal forebrain subcortical projections. Brain Struct. Funct. 224, 1097–1117 (2019).

8. Do, J. P. et al. Cell type-specific long-range connections of basal forebrain circuit. Elife 5, 1–17 (2016).

9. Maness, E. B. et al. Role of the locus coeruleus and basal forebrain in arousal and attention. Brain Res. Bull. 188, 47–58 (2022).

10. Kim, T. et al. Cortically projecting basal forebrain parvalbumin neurons regulate cortical gamma band oscillations. Proc. Natl. Acad. Sci. 112, 3535–3540 (2015).

11. Xu, M. et al. Basal forebrain circuit for sleep-wake control. Nat. Neurosci. 18, 1641–1647 (2015).

12. McKenna, J. T. et al. Basal Forebrain Parvalbumin Neurons Mediate Arousals from Sleep Induced by Hypercarbia or Auditory Stimuli. Curr. Biol. 30, 2379–2385.e4 (2020).

13. Etter, G. et al. Optogenetic gamma stimulation rescues memory impairments in an Alzheimer’s disease mouse model. Nat. Commun. 10, 1–11 (2019).

14. Burk, J. . & Sarter, M. Dissociation between the attentional functions mediated via basal forebrain cholinergic and GABAergic neurons. Neuroscience 105, 899–909 (2001).

15. McGaughy, J., Dalley, J. W., Morrison, C. H., Everitt, B. J. & Robbins, T. W. Selective Behavioral and Neurochemical Effects of Cholinergic Lesions Produced by Intrabasalis Infusions of 192 IgG-Saporin on Attentional Performance in a Five-Choice Serial Reaction Time Task. J. Neurosci. 22, 1905–1913 (2002).

16. Roßner, S. Cholinergic immunolesions by 192IgG-saporin—a useful tool to simulate pathogenic aspects of alzheimer’s disease. Int. J. Dev. Neurosci. 15, 835–850 (1997).

17. Muir, J. L., Page, K. J., Sirinathsinghji, D. J. S., Robbins, T. W. & Everitt, B. J. Excitotoxic lesions of basal forebrain cholinergic neurons: Effects on learning, memory and attention. Behav. Brain Res. 57, 123–131 (1993).

18. Roland, J. J., Janke, K. L., Servatius, R. J. & Pang, K. C. H. GABAergic neurons in the medial septum-diagonal band of Broca (MSDB) are important for acquisition of the classically conditioned eyeblink response. Brain Struct. Funct. 219, 1231–7 (2014).

19. Lin, S.-C. & Nicolelis, M. a L. Neuronal ensemble bursting in the basal forebrain encodes salience irrespective of valence. Neuron 59, 138–49 (2008).

20. Avila, I. & Lin, S.-C. Motivational Salience Signal in the Basal Forebrain Is Coupled with Faster and More Precise Decision Speed. PLoS Biol. 12, e1001811 (2014).

21. Hangya, B., Ranade, S. P., Lorenc, M. & Kepecs, A. Central Cholinergic Neurons Are Rapidly Recruited by Reinforcement Feedback. Cell 162, 1155–1168 (2015).

22. Hegedüs, P., Velencei, A., Belval, C.-H. de, Heckenast, J. & Hangya, B. Training protocol for probabilistic Pavlovian conditioning in mice using an open-source head-fixed setup. STAR Protoc. 2, 100795 (2021).

23. Trask, S. & Helmstetter, F. J. Unique roles for the anterior and posterior retrosplenial cortices in encoding and retrieval of memory for context. Cereb. Cortex 32, 3602–3610 (2022).

24. Stepanichev, M., Lazareva, N., Tukhbatova, G., Salozhin, S. & Gulyaeva, N. Transient disturbances in contextual fear memory induced by Aβ(25-35) in rats are accompanied by cholinergic dysfunction. Behav. Brain Res. 259, 152–157 (2014).

25. Tsai, T. C., Yu, T. H., Hung, Y. C., Fong, L. I. & Hsu, K. Sen. Distinct Contribution of Granular and Agranular Subdivisions of the Retrosplenial Cortex to Remote Contextual Fear Memory Retrieval. J. Neurosci. 42, 877–893 (2022).

26. Mitsushima, D., Sano, A. & Takahashi, T. A cholinergic trigger drives learning-induced plasticity at hippocampal synapses. Nat. Commun. 4, 1–10 (2013).

27. Pan, T. T. et al. Retrosplenial Cortex Effects Contextual Fear Formation Relying on Dysgranular Constituent in Rats. Front. Neurosci. 16, 1–10 (2022).

28. Tronson, N. C. et al. Segregated populations of hippocampal principal CA1 neurons mediating conditioning and extinction of contextual fear. J. Neurosci. 29, 3387–3394 (2009).

29. Kocsis, B. et al. Huygens synchronization of medial septal pacemaker neurons generates hippocampal theta oscillation. Cell Rep. 40, 111149 (2022).

30. Sturgill, J. F. et al. Basal forebrain-derived acetylcholine encodes valence-free reinforcement prediction error. bioRxiv (2020) doi:10.1101/2020.02.17.953141.

31. Hegedüs, P., Sviatkó, K., Király, B., Martínez-Bellver, S. & Hangya, B. Cholinergic activity reflects reward expectations and predicts behavioral responses. iScience 26, 105814 (2023).

32. Szőnyi, A. et al. Median raphe controls acquisition of negative experience in the mouse. Science 366, eaay8746 (2019).

33. Sos, K. E. et al. Cellular architecture and transmitter phenotypes of neurons of the mouse median raphe region. Brain Struct. Funct. 222, 287–299 (2017).

34. Varga, V. et al. Fast synaptic subcortical control of hippocampal circuits. Science (80-. ). 326, (2009).

35. Freund, T. F. & Antal, M. GABA-containing neurons in the septum control inhibitory interneurons in the hippocampus. Nature 336, 403–405 (1988).

36. McKenna, J. T. et al. Distribution and intrinsic membrane properties of basal forebrain GABAergic and parvalbumin neurons in the mouse. J. Comp. Neurol. 521, 1225–1250 (2013).

37. Freund, T. F. GABAergic septohippocampal neurons contain parvalbumin. Brain Res. 478, 375– 381 (1989).

38. Tian, J. et al. Dissection of the long-range projections of specific neurons at the synaptic level in the whole mouse brain. Proc. Natl. Acad. Sci. 119, 1–10 (2022).

39. Wrenn, C. C. & Wiley, R. G. The behavioral functions of the cholinergic basalforebrain : lessons from 192 IgG-SAPORIN. Int. J. Dev. Neurosci. 16, 595–602 (1998).

40. McGaughy, J., Everitt, B. J., Robbins, T. W. & Sarter, M. The role of cortical cholinergic afferent projections in cognition: impact of new selective immunotoxins. Behav. Brain Res. 115, 251– 63 (2000).

41. Frick, K. M., Kim, J. J. & Baxter, M. G. Effects of complete immunotoxin lesions of the cholinergic basal forebrain on fear conditioning and spatial learning. Hippocampus 14, 244–54 (2004).

42. Calandreau, L., Jaffard, R. & Desmedt, A. Dissociated roles for the lateral and medial septum in elemental and contextual fear conditioning. Learn. Mem. 14, 422–429 (2007).

43. Dwyer, T. A., Servatius, R. J. & Pang, K. C. H. Noncholinergic Lesions of the Medial Septum Impair Sequential Learning of Different Spatial Locations. J. Neurosci. 27, 299–303 (2007).

44. Laszlovszky, T. et al. Distinct synchronization, cortical coupling and behavioral function of two basal forebrain cholinergic neuron types. Nat. Neurosci. 23, 992–1003 (2020).

45. Ang, S. T. et al. GABAergic neurons of the medial septum play a nodal role in facilitation of nociception-induced affect. Sci. Rep. 5, 15419 (2015).

46. Kai, Y., Oomura, Y. & Shimizu, N. Responses of rat lateral hypothalamic neurons to periaqueductal gray stimulation and nociceptive stimuli. Brain Res. 461, 107–117 (1988).

47. Berthoud, H.-R. & Münzberg, H. The lateral hypothalamus as integrator of metabolic and environmental needs: From electrical self-stimulation to opto-genetics. Physiol. Behav. 104, 29–39 (2011).

48. Burstein, R., Cliffer, K. & Giesler, G. Direct somatosensory projections from the spinal cord to the hypothalamus and telencephalon. J. Neurosci. 7, 4159–4164 (1987).

49. Hegedüs, P., Heckenast, J. & Hangya, B. Differential recruitment of ventral pallidal e-types by behaviorally salient stimuli during Pavlovian conditioning. iScience 24, 102377 (2021).

50. Zhang, G.-W. et al. A Non-canonical Reticular-Limbic Central Auditory Pathway via Medial Septum Contributes to Fear Conditioning. Neuron 97, 406–417.e4 (2018).

51. Cohen, J. Y., Amoroso, M. W. & Uchida, N. Serotonergic neurons signal reward and punishment on multiple timescales. Elife 4, 06346 (2015).

52. Ranade, S. P. & Mainen, Z. F. Transient firing of dorsal raphe neurons encodes diverse and specific sensory, motor, and reward events. J. Neurophysiol. 102, 3026–37 (2009).

53. Sheehan, T. P., Chambers, R. A. & Russell, D. S. Regulation of affect by the lateral septum: implications for neuropsychiatry. Brain Res. Rev. 46, 71–117 (2004).

54. Rocamora, N. et al. Expression of NGF and NT3 mRNAs in Hippocampal Interneurons Innervated by the GABAergic Septohippocampal Pathway. J. Neurosci. 16, 3991–4004 (1996).

55. Solari, N. & Hangya, B. Cholinergic modulation of spatial learning, memory and navigation. Eur. J. Neurosci. 48, 2199–2230 (2018).

56. Miller, A. M. P., Vedder, L. C., Law, L. M. & Smith, D. M. Cues, context, and long-term memory: the role of the retrosplenial cortex in spatial cognition. Front. Hum. Neurosci. 8, 1–15 (2014).

57. Borhegyi, Z., Varga, V., Szilagyi, N., Fabo, D. & Freund, T. F. Phase segregation of medial septal GABAergic neurons during hippocampal theta activity. J. Neurosci. 24, 8470–8479 (2004).

58. Villar, P. S., Hu, R. & Araneda, R. C. Long-Range GABAergic Inhibition Modulates Spatiotemporal Dynamics of the Output Neurons in the Olfactory Bulb. J. Neurosci. 41, 3610– 3621 (2021).

59. Gracia-Llanes, F. J. et al. GABAergic basal forebrain afferents innervate selectively GABAergic targets in the main olfactory bulb. Neuroscience 170, 913–922 (2010).

60. Salib, M. et al. GABAergic Medial Septal Neurons with Low-Rhythmic Firing Innervating the Dentate Gyrus and Hippocampal Area CA3. J. Neurosci. 39, 4527–4549 (2019).

61. Viney, T. J. et al. Shared rhythmic subcortical GABAergic input to the entorhinal cortex and presubiculum. Elife 7, 1–35 (2018).

62. McDonald, A. J., Muller, J. F. & Mascagni, F. Postsynaptic targets of GABAergic basal forebrain projections to the basolateral amygdala. Neuroscience 183, 144–159 (2011).

63. Jiang, L. et al. Cholinergic Signaling Controls Conditioned Fear Behaviors and Enhances Plasticity of Cortical-Amygdala Circuits. Neuron 90, 1057–1070 (2016).

64. Záborszky, L. et al. Specific Basal Forebrain–Cortical Cholinergic Circuits Coordinate Cognitive Operations. J. Neurosci. 38, 9446–9458 (2018).

65. Staib, J. M., Della Valle, R. & Knox, D. K. Disruption of medial septum and diagonal bands of Broca cholinergic projections to the ventral hippocampus disrupt auditory fear memory. Neurobiol. Learn. Mem. 152, 71–79 (2018).

66. Marks, W. D., Yokose, J., Kitamura, T. & Ogawa, S. K. Neuronal Ensembles Organize Activity to Generate Contextual Memory. Front. Behav. Neurosci. 16, 1–16 (2022).

67. Stephenson-Jones, M. et al. Opposing Contributions of GABAergic and Glutamatergic Ventral Pallidal Neurons to Motivational Behaviors. Neuron 105, 921–933.e5 (2020).

68. Wulff, A. B., Tooley, J., Marconi, L. J. & Creed, M. C. Ventral pallidal modulation of aversion processing. Brain Res. 1713, 62–69 (2019).

69. Farrell, M. R. et al. Ventral Pallidum GABA Neurons Mediate Motivation Underlying Risky Choice. J. Neurosci. 41, 4500–4513 (2021).

70. Fournier, D. I., Monasch, R. R., Bucci, D. J. & Todd, T. P. Retrosplenial cortex damage impairs unimodal sensory preconditioning. Behav. Neurosci. 134, 198–207 (2020).

71. Xu, M. et al. Basal forebrain circuit for sleep-wake control. Nat. Neurosci. 18, 1641–1647 (2015).

72. Lozano-Montes, L. et al. Optogenetic Stimulation of Basal Forebrain Parvalbumin Neurons Activates the Default Mode Network and Associated Behaviors. Cell Rep. 33, 108359 (2020).

73. Aguilar, D. D. & McNally, J. M. Subcortical control of the default mode network: Role of the basal forebrain and implications for neuropsychiatric disorders. Brain Res. Bull. 185, 129–139 (2022).

74. Lewthwaite, R. & Wulf, G. Optimizing motivation and attention for motor performance and learning. Curr. Opin. Psychol. 16, 38–42 (2017).

75. Kvitsiani, D. et al. Distinct behavioural and network correlates of two interneuron types in prefrontal cortex. Nature 498, 363–6 (2013).

76. Pi, H.-J. et al. Cortical interneurons that specialize in disinhibitory control. Nature 503, 521– 524 (2013).

77. Solari, N., Sviatkó, K., Laszlovszky, T., Hegedüs, P. & Hangya, B. Open Source Tools for Temporally Controlled Rodent Behavior Suitable for Electrophysiology and Optogenetic Manipulations. Front. Syst. Neurosci. 12, 18 (2018).

78. Janssen, P. & Shadlen, M. N. A representation of the hazard rate of elapsed time in macaque area LIP. Nat. Neurosci. 8, 234–41 (2005).

79. Ralls, K. Auditory sensitivity in mice: Peromyscus and Mus musculus. Anim. Behav. 15, 123– 128 (1967).

80. Najafi, F., Giovannucci, A., Wang, S. S. H. & Medina, J. F. Coding of stimulus strength via analog calcium signals in Purkinje cell dendrites of awake mice. Elife 3, e03663 (2014).

81. Ferguson, J. E., Boldt, C. & Redish, A. D. Creating low-impedance tetrodes by electroplating with additives. Sensors Actuators A Phys. 156, 388–393 (2009).

82. Széll, A., Martínez-Bellver, S., Hegedüs, P. & Hangya, B. OPETH: Open Source Solution for Real- Time Peri-Event Time Histogram Based on Open Ephys. Front. Neuroinform. 14, 1–19 (2020).

83. Endres, D. M. & Schindelin, J. E. A New Metric for Probability Distributions. IEEE Trans. Inf. Theory 49, 1858–1860 (2003).

84. Fraser, G. W. & Schwartz, A. B. Recording from the same neurons chronically in motor cortex. J. Neurophysiol. 107, 1970–1978 (2012).

85. Paxinos, G. & Franklin, K. B. J. The Mouse Brain in Stereotaxic Coordinates, 2nd edition. Academic Press (2001).

86. Cohen, J. Y., Haesler, S., Vong, L., Lowell, B. B. & Uchida, N. Neuron-type-specific signals for reward and punishment in the ventral tegmental area. Nature 482, 85–8 (2012).

87. Schmitzer-Torbert, N., Jackson, J., Henze, D., Harris, K. & Redish, a D. Quantitative measures of cluster quality for use in extracellular recordings. Neuroscience 131, 1–11 (2005).

88. Royer, S. et al. Control of timing, rate and bursts of hippocampal place cells by dendritic and somatic inhibition. Nat. Neurosci. 15, 769–75 (2012).

89. Fujisawa, S., Amarasingham, A., Harrison, M. T. & Buzsáki, G. Behavior-dependent short-term assembly dynamics in the medial prefrontal cortex. Nat. Neurosci. 11, 823–33 (2008).

90. Lerner, T. N. et al. Intact-Brain Analyses Reveal Distinct Information Carried by SNc Dopamine Subcircuits. Cell 162, 635–647 (2015).

91. Fox, S. R. & Deneris, E. S. Engrailed is required in maturing serotonin neurons to regulate the cytoarchitecture and survival of the dorsal raphe nucleus. J. Neurosci. 32, 7832–7842 (2012).

